# Phased grapevine genome sequence of an *Rpv12* carrier for biotechnological exploration of resistance to *Plasmopara viticola*

**DOI:** 10.1101/2022.08.06.503030

**Authors:** Bianca Frommer, Sophia Müllner, Daniela Holtgräwe, Prisca Viehöver, Bruno Hüttel, Reinhard Töpfer, Bernd Weisshaar, Eva Zyprian

**Author notes:** **Correspondence:** Bernd Weisshaar, Reinhard Töpfer. These authors contributed equally to this work and share first authorship (Bianca Frommer, Sophia Müllner). Email addresses: BF SM DH PV BH RT BW EZ.

## Abstract

The downy mildew disease caused by the oomycete *Plasmopara viticola* is a serious threat for grapevine and can cause enormous yield losses in viticulture. The quantitative trait locus *Rpv12,* mediating resistance against *P. viticola*, was originally found in Asian *Vitis amurensis*. This locus and its genes were analyzed here in detail. A haplotype-separated genome sequence of the diploid *Rpv12*-carrier Gf.99-03 was created and annotated. The defense response against *P. viticola* was investigated in an infection time-course RNA-Seq experiment, revealing approximately 600 up-regulated *Vitis* genes during host-pathogen interaction. The *Rpv12* regions of the resistance conferring and the sensitivity encoding Gf.99-03 haplotypes were structurally and functionally compared to each other. Two different clusters of resistance-related genes were identified within the *Rpv12* locus. One cluster carries a set of four differentially expressed genes with three *ACCELERATED CELL DEATH 6*-like genes. The other cluster carries a set of six resistance gene analogues related to qualitative pathogen resistance. The *Rpv12* locus and its candidate genes for *P. viticola* resistance provide a precious genetic resource for *P. viticola* resistance breeding. Newly developed co-segregating simple sequence repeat markers in close proximity to the *R*-genes enable its improved applicability in marker-assisted grapevine breeding.

## 1 Introduction

With the transatlantic migration of people, various plant pathogens were introduced to Europe during the 19th century. These include the causative agent of downy mildew: *Plasmopara viticola* ((Berk. & Curt.) Berl. & de Toni), an obligate biotrophic oomycete, member of the *Peronosporales* (Gessler et al., 2011). Already in the 19th century, breeding programs were started to delimit the damage caused by downy mildew to viticulture (Bavaresco, 2018). These attempted to introgress resistance traits present in American or Asian *Vitis* species into European grapevine varieties, enabled by their general diploidity and cross fertility. First results were discouraging because of the prominent fox tones co-inherited from the resistant *Vitis* accessions (Töpfer et al., 2011; Reynolds, 2022). Later, after several generations of back crosses to European *V. vinifera* noble varieties, newly bred resistant varieties such as ‘Regent’ (1967) and ‘Johannite’ (1968) were introduced to German viticulture (https://www.vivc.de/) (Statistisches Bundesamt (Destatis), 2016).

However, these grapevine varieties carry only one resistance locus to each downy and powdery mildew and may become susceptible to newly emerging pathogen strains (Kast, 2001; Peressotti et al., 2010; Heyman et al., 2021). Genetic analysis during the last decades revealed various resistance loci from several *Vitis* sources. Using marker assisted selection these can be combined to establish improved durability of the resistance trait (Eibach et al., 2007; Consortium, 2016). To date, more than 30 different quantitative trait loci (QTLs) associated with *P. viticola* resistance are described (https://www.vivc.de/). The most employed ones comprise different alleles of *Rpv3* (e. g. *Rpv3.1* and *Rpv3.3*), *Rpv10* and *Rpv12* (*Rpv* for Resistance to *Plasmopara viticola*) (Welter et al., 2007; Di Gaspero et al., 2012; Schwander et al., 2012; Venuti et al., 2013). In general, resistance loci used in breeding are associated with hypersensitive responses (HR), a defense mechanism operated through local necrosis through the topical production of reactive oxygen species (ROS) (Possamai et al., 2020).

The QTL *Rpv12* is located on chr14 and was identified in several independent introgressions of *V. amurensis*. It is present in the genomes of diverse varieties like ‘Michurinetś, ‘Zarja severá, as well as in ‘Kunbarat’, ‘Petrá, ‘Lelá, ‘Kunleaný and ‘Milá (Venuti et al., 2013). Recently, it was shown that *Rpv12* was transmitted by *V. amurensis* to the grapevine variety ‘Merlot Khoruś based on introgression maps (Foria et al., 2022). The locus *Rpv12* is delimited by the simple sequence repeat (SSR) markers UDV-350 and UDV-370 (Venuti et al., 2013). The corresponding region of the grapevine reference genome sequence (12X.v2), which is derived from the susceptible genotype PN40024 (Jaillon et al., 2007; Canaguier et al., 2017), covers twelve typical resistance gene analogues (RGAs) of the CNL Type (CC-NBS-LRR, coiled coil, nucleotide binding site and leucine rich repeat encoding genes (see (Han, 2019)).

To investigate the mechanisms of *P. viticola* resistance mediated by *Rpv12*, both haplotypes of a heterozygous *Rpv12*-carrying grapevine genotype were sequenced and separated. The haplotype sequences were analyzed for candidate resistance genes present specifically at the *Rpv12* locus (positional candidates). In addition, a time-resolved gene expression study after *P. viticola* inoculation revealed gene activities leading to successful plant defense (functional candidates).

## 2 Results

### 2.1 Pedigree of Gf.99-03

The trio binning approach (Koren et al., 2018) was employed to generate fully phased haplotype sequences. This requires sequence data from both parental genotypes and their offspring. Unfortunately, one or both parental genotypes of well-known *Rpv12*-carriers like ‘Kunbarat’ and ‘Michurinetś are unavailable. For this reason, the parent-child trio 65-153-18 (*Rpv12*-carrier), Gf.43- 21 (susceptible) and Gf.99-03 (*Rpv12*-carrier) was chosen for analysis. The pedigree (Figure 1) was verified by 154 segregating SSR markers (Table S1). In addition, its relationship to the ancestors ‘Blaufraenkisch’, ‘Calandró, ‘Regent’ and ‘Dominá was confirmed using 83 markers (Table S2).

**Figure 1.**
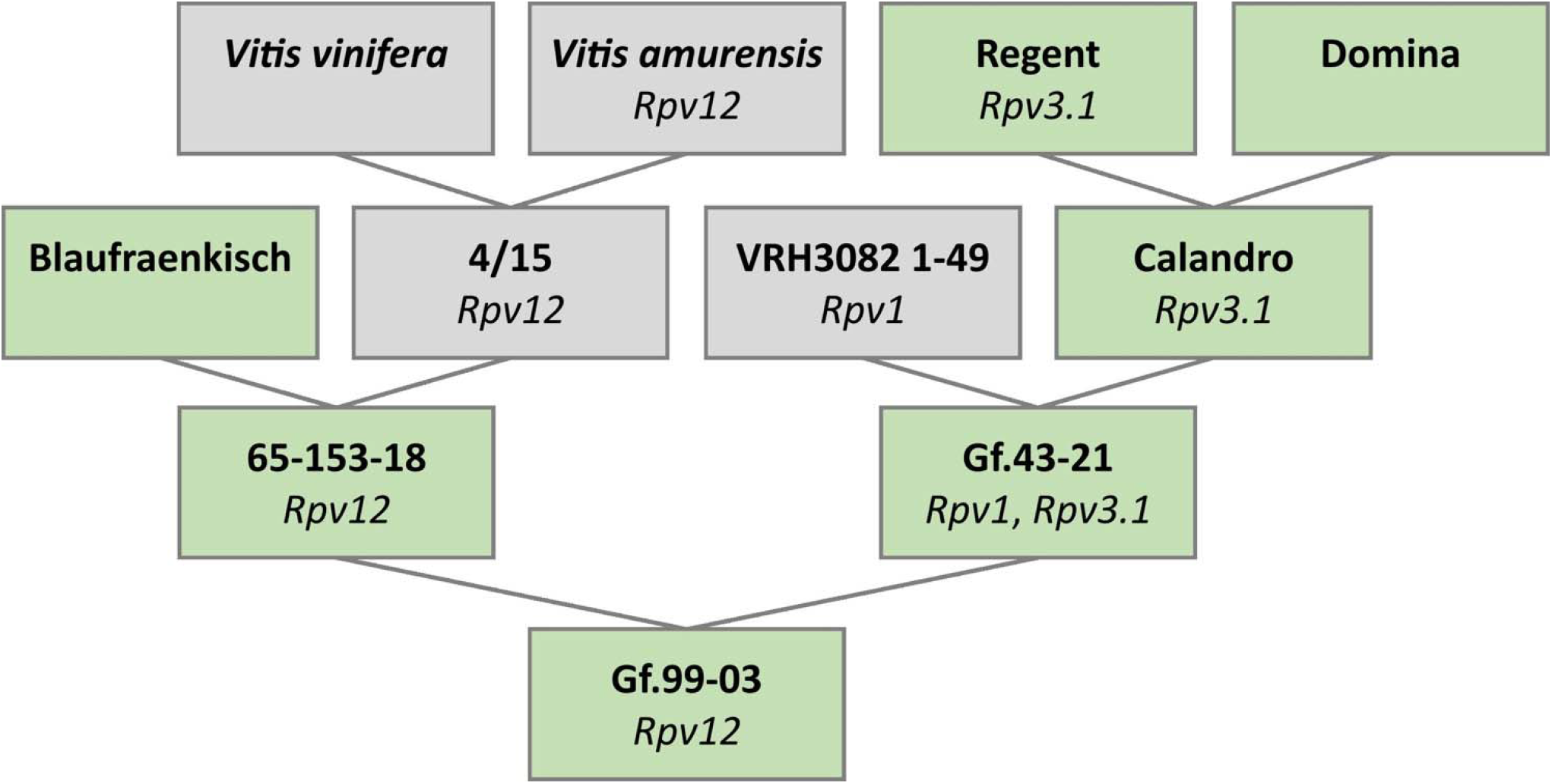
Pedigree of the heterozygous *Rpv12*-carrier Gf.99-03 chosen for phased genome sequence assembly. This pedigree shows the confirmed relationship between the genotypes 65-153- 18, Gf.43-21 and Gf.99-03. Genotypes in green were available while genotypes in grey were not available. Identified resistance-associated loci are given below the genotype.

The pedigree of Gf.43-21 was not completely clarified, due to unavailability of the genotype VRH3082 1-49 (Figure 1). However, ‘Calandró (Rpv3.1, ‘Regent’ x ‘Dominá) as the second ancestor was confirmed. Since this ancestry could potentially transmit the resistance loci *Rpv1* and *Rpv3.1*, their absence in the *Rpv12*-carrier Gf.99-03 was checked and verified by analysis of markers linked to these two loci (Sc35_2 (GenBank no. GF111546.1), Sc34_8 (GenBank no. GF111545.1), GF18-06 (Schwander et al., 2012) and GF18-08 (Zyprian et al., 2016).

### 2.2 *Plasmopara viticola* interaction with *Rpv12* positive and *Rpv12* negative genotypes

To characterize the resistance mechanism caused by *Rpv12*, the development of *P. viticola* sporangiophores and local necrotic tissue reactions indicative of HR were investigated on genotypes Gf.99-03, Gf.43-21, 65-153-18 and ‘Italiá (Figure 2 (A-D)). While ‘Italiá as susceptible control allowed the formation of many sporangiophores, the *Rpv*-carriers 65-153-18 (*Rpv12*) and Gf.43-21 (*Rpv1*, *Rpv3.1*) restricted the development to none or only few sporangiophores within five days post infection (dpi), as did their offspring Gf.99-03 (*Rpv12*). Gf.99-03 exhibited HR as indicated by small necrotic lesions at the infection site (Figure 2 (D)). Defense responses of Gf.99-03 resulted in ROS formation (Figure 2 (F); Figure S1). Remarkably, *P. viticola* formed smaller, but conspicuously more haustoria in the *Rpv12* carrier leaf than in the susceptible ‘Italiá (Figure 2 (G, H)).

**Figure 2.**
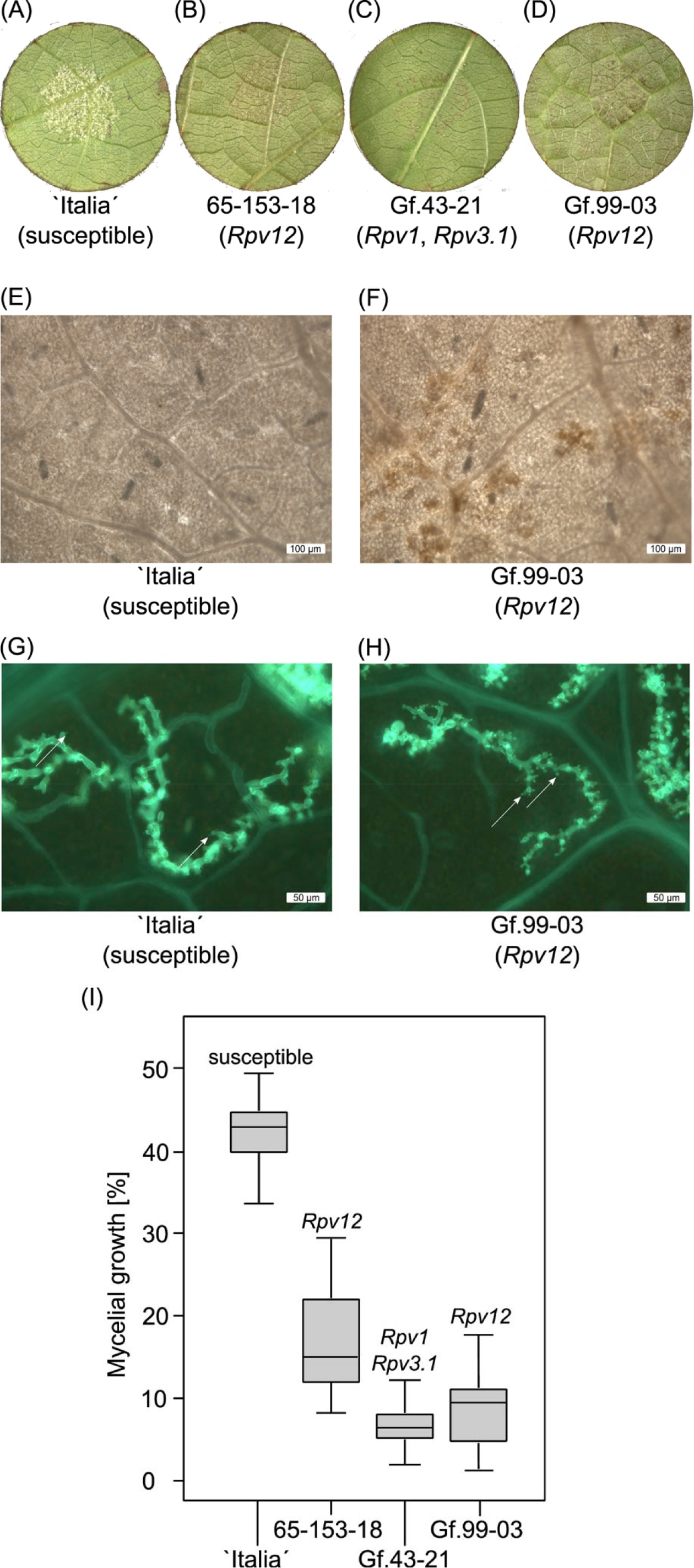
P*l*asmopara *viticola* infection of *Rpv12*-carrying and non *Rpv12*-carrying genotypes. **(A-D)** Leaf disc assay of different genotypes 5 dpi after artificial inoculation with *P. viticola*. **(A)** Leaf discs from genotype ‘Italiá covered by sporangiophores. **(B)** 65-153-18 (*Rpv12*) shows strongly reduced sporangiophore development and HR visible as necrotic spots. **(C)** Genotype Gf.43-21 (*Rpv1*, *Rpv3.1*) exhibits no sporangiophores but HR visible as necrotic spots. **(D)** Gf.99-03 (*Rpv12*) has very little sporangiophores and HR is visible as necrotic spots. **(E, F)** Nitrotetrazolium blue chloride staining for detection of ROS. *P. viticola* infected leaf discs of ‘Italiá (susceptible) and Gf.99-03 (*Rpv12*) were incubated for 24 h and were stained with Nitrotetrazolium blue chloride. **(E)** The susceptible genotype ‘Italiá shows no ROS. **(F)** Genotype Gf.99-03 (*Rpv12*) produced ROS that led to visible brown lesions. **(G, H)** Comparison of haustoria formation. Fluorescence microscopic images of *P. viticola* infestation on the genotypes ‘Italiá and Gf.99-03 at 72 hpi. Mycelium was stained with alkaline aniline blue. Arrows are pointing on haustoria. **(G)** ‘Italiá (susceptible) shows less haustoria and a widely spread mycelium. **(H)** Gf.99-03 (*Rpv12*) shows many small haustoria with reduced width. **(I)** Comparison of mycelial growth. Mycelial density was evaluated on leaves of Gf.99-03 and its parental genotypes Gf.43-21 and 65-153-18 including ‘Italiá as susceptible control. The results are visualized as box-plots with standard deviation (Welch’s t-test). Relevant resistance loci are given above the error bar. The median is marked as a thick black line within the box while the whiskers subscript the minimum and maximum of all data. The data of the variety ‘Italiá as reference was already published (Müllner and Zyprian, 2022).

Infected leaves were investigated to compare the progress of mycelial development in the mesophyll (Figure 2 (I)). The presence of the *Rpv12* locus in Gf.99-03 and its paternal genotype 65-153-18, reduced the mycelial growth to half. The maternal genotype Gf.43-21, which carries a combination of two different *Rpv* resistance loci, showed a stronger reduction of mycelial development than 65-153-18. Astonishingly, Gf.99-03 showed reduced mycelial growth compared to 65-153-18, even though it carries only *Rpv12* as known resistance locus against *P. viticola*.

### 2.3 Sequence data and phase-separated genome sequence assembly of Gf.99-03

The Gf.99-03 long reads consisted of ∼71 Gbp (average read length 11,132 bp; N50 15,915 bp) of data. Assuming a size of ∼500 Mbp for each haplotype, the genome was covered at 1n 142-fold or ∼70-fold for each haplotype. Illumina short read data of ∼116 Gbp (maternal genotype Gf.43-21) and ∼144 Gbp (paternal genotype 65-153-18) were used, providing more than 200-fold genome coverage each.

The long reads were divided into the Gf.43-21 read bin containing 46.61 % of all bases, the 65-153- 18 bin comprising 51.14 % of all bases, and the unassignable read bin containing 2.25 % of all bases (Table S3). As expected for successful binning, the parental bins were each holding about half of the total bases. The unassigned bin comprised only a small fraction of all reads and contained mostly short sequences (avg. length 3,638 bp).

The parental bins were assembled separately, yielding two haplotype assemblies that were designated Gf9921 for the maternal haplotype derived from Gf.43-21 and Gf9918 for the paternal haplotype derived from 65-153-18 (Table 1). 90% of the sequence information of both assemblies is represented by less than 240 contigs. The additional ∼1,000 contigs in each of the assemblies are mostly relatively short, presumably caused by high content of repetitive sequences.

**Table 1.** Statistics of the Gf.99-03 assembly. Gf9921 denotes the maternal haplotype assembly and Gf9918 the paternal haplotype assembly, both derived from phase-separated reads of Gf.99-03.

|  | **Gf9921** | **Gf9918** | **Gf.99-03 assembly** |
| --- | --- | --- | --- |
| **contigs** | 1,245 | 1,253 | 2,498 |
| **size [bp]** | 511,095,651 | 544,110,372 | 1,055,206,023 |
| **largest contig [bp]** | 14,810,751 | 18,073,517 | 18,073,517 |
| **N50 length [bp]** | 3,294,480 | 3,920,956 | 3,763,673 |
| **N50 contigs** | 42 | 38 | 79 |
| **N90 length [bp]** | 200,578 | 203,624 | 201,530 |
| **N90 contigs** | 239 | 235 | 474 |

The contigs of both haplotypes were assigned to two sets of (homologous) pseudochromosomes based on reciprocal best BLAST hits (RBHs) and synteny with other *Vitis* genome sequence assemblies (Table S4). For Gf9921, 237 contigs representing 85.64 % of all bases, and for Gf9918 195 contigs representing 83.10 % of all bases were ordered into pseudochromosomes (Figure S2, Figure S3).

In general, the lengths of the final homologous pseudochromosomes of Gf9921 and Gf9918 were similar (Figure 3 (A), Table S5). Only the homologs of chr10, chr13 and chr19 differed in their length by more than 3 Mbp. However, the artificial pseudochromosome chrNh with contigs that had no RBH with any of the syntenic genome sequences (see Experimental Procedures) was significantly larger by ∼18.5 Mbp for the Gf9918 assembly as compared to Gf9921. The total size of the Gf9918 assembly is around 33 Mbp larger than that of Gf9921.

**Figure 3.**
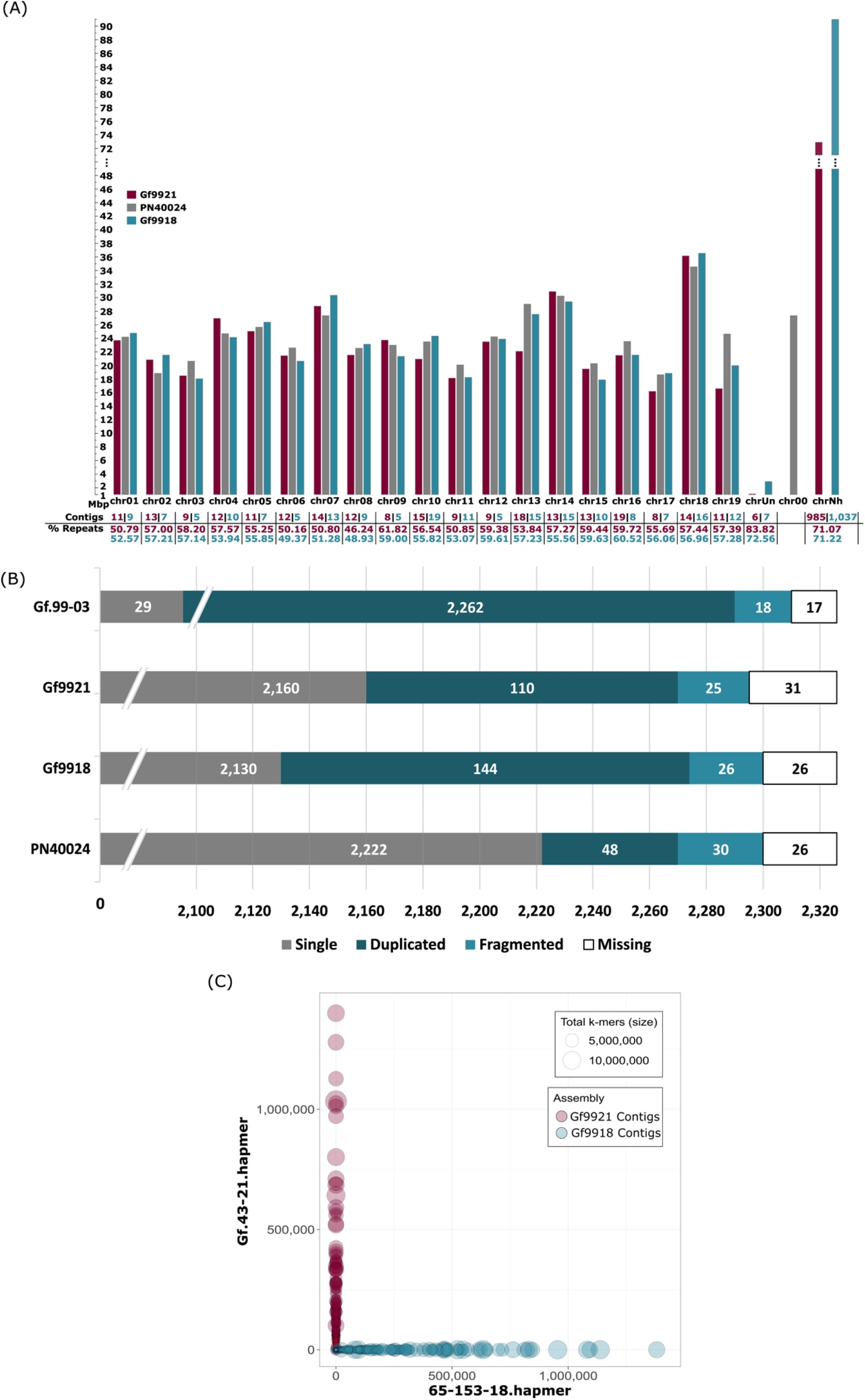
Pseudochromosome lengths, BUSCO analysis of the Gf.99-03 haplotypes and of PN40024 and bob plot of the Gf.99-03 contigs. **(A)** Pseudochromosome lengths of both Gf.99-03 haplotypes and of PN40024 12X.v2. The lengths and values are shown in magenta for Gf9921, in turquoise for Gf9918, and in grey for PN40024. chrUn, chr00 and chrNh represent artificial contig collections. The reference sequence PN40024 12X.v2 contains a pseudochromosome chr00 for sequences not assigned to chromosomes, this collection has been created as well and is referred to here as “chrUn” for Gf9921 and Gf9918. chrNh collects contigs that cannot be assigned to chromosomal locations based on similarity to established *Vitis* genome sequences. The scale on the y-axis and the lengths of chrNh are interrupted at 48 Mbp. **(B)** Number of plant core genes for Gf.99- 03 and PN40024.2,326 plant core genes were searched with BUSCO. Note that the bar graph is truncated at the left and focuses on only duplicated, fragmented and missing BUSCOs. The track Gf.99-03 combines the summed-up results of the single haplotypes. **(C)** Contigs are displayed as circles. Magenta circles represent Gf9921 contigs and turquoise circles Gf9918 contigs. The circle size depends on the total number of k-mers assigned to a given contig. The axes indicate the number of k-mers found in the hap-mer sets (y-axis Gf.43-21 hap-mers, x-axis 65-153-18 hap-mers; the analysis tool Mercury refers to haplotype-specific k-mers as hap-mers).

To validate the completeness of the haplotype assemblies, the presence of plant core genes was determined with Benchmarking Universal Single-Copy Orthologs (BUSCO) (Figure 3 (B)). Despite a very low portion of missing and fragmented plant core genes for the combined assembly (0.9 % fragmented, 0.6 % missing) and for each haplotype, the content of duplicated genes is rather high for the haplotypes (4.7 % Gf9921, 6.2 % Gf9918), compared to the reference PN40024 with 2.1 % duplications.

Analysis of the Gf.99-03 long reads and of the haplotype assemblies revealed no significant contaminations. More than 99 % of all database hits were hits against *Vitis* sequences. The few other hits were mostly hits against the tree *Spondias tuberosa* or against sequences of the genera *Berberidopsis* and *Ampelopsis*.

### 2.4 Phasing status and k-mer completeness

For verification of the phasing status, more than 470 SSR marker candidates were selected from literature. About 250 could be assigned to a pseudochromosome position without mismatches in the assay primer sequences. From these ∼250 markers, 203 amplification products were generated and tested for inheritance using DNA of Gf.99-03 and its parents.

Out of these 203 markers, 154 segregating and 22 non-segregating markers were identified for Gf.99-03 (had an appropriate PCR product; PCR product matched to one of the parental haplotypes) and 27 had to be discarded (no or multiple products). Every segregating marker tested (154) supported a full phasing (Table S1), e. g. no haplotype switching was identified.

The k-mer based phasing assessment analysis with Merqury (Rhie et al., 2020) resulted in a base level quality value (QV) of 35.44 and a k-mer completeness of >97 % for the Gf.99-03 genome assembly (Table S6). The k-mer completeness for the haplotypes was estimated to >99 %. The analyses indicated a higher content of heterozygous sequence (red 1-copy peak) than of homozygous sequence (blue 2-copy peak), almost no missing sequences (read-only peak) and a few artificial duplications (green peak) (Figure S4). The k-mer switch error rate was <0.03 % per haplotype by only allowing 10 switches per 20 kbp (Table S6). The blob plot showed haplotype-pure contigs (no mixtures between haplotypes, Figure 3 (C)).

### 2.5 Gene annotation assisted by RNA-Seq data and extraction of RGAs

Annotation of both haplotype assemblies was supported by RNA-Seq data generated from leaf, tendril, root and stem of Gf.99-03 (Table S7). Gene annotation (see Experimental Procedures) resulted in detection of 34,713 protein-coding genes for Gf9921 and 36,290 protein-coding genes for Gf9918. In addition, 1,070 and 855 tRNA coding genes were predicted, respectively.

Since this investigation aims to investigate the resistance mediated by *Rpv12*, special focus was laid on resistance gene analogues. Using the RGA annotation pipeline, 2,288 Gf9921 genes and 2,268 Gf9918 genes were classified as RGA (Table S8, Figure S6). Relatively similar numbers of RGAs were identified for both haplotypes over the RGA classes, however the CNL genes, TIR-X (Toll/Interleukin-1 Receptor like with unknown domain) genes and RLP (Receptor Like Protein) genes were more numerous in the *Rpv12* carrying haplotype Gf9918 than in the susceptible haplotype Gf9921.

### 2.6 Comparative RNA-Seq analysis of a *Plasmopara viticola* infection time course experiment challenging grapevine

Based on phenotypic analyses of the infection process, time points and duration for an infection time course were selected. For five time points (0 hours post inoculation (hpi), 6 hpi, 12 hpi, 24 hpi, 48 hpi) RNA was extracted and sequenced (Table S9).

A Principal Component Analysis (PCA) of the RNA-Seq read counts showed time-separated data points in the time course order (x-axis) and condition-separated data points along the y-axis (Figure 4 (A)). The data points of the infection start time at 0 hpi were not separated by condition, but clustered together. In addition, the triplicates of each time point and condition clustered together, indicating high experimental reproducibility.

**Figure 4.**
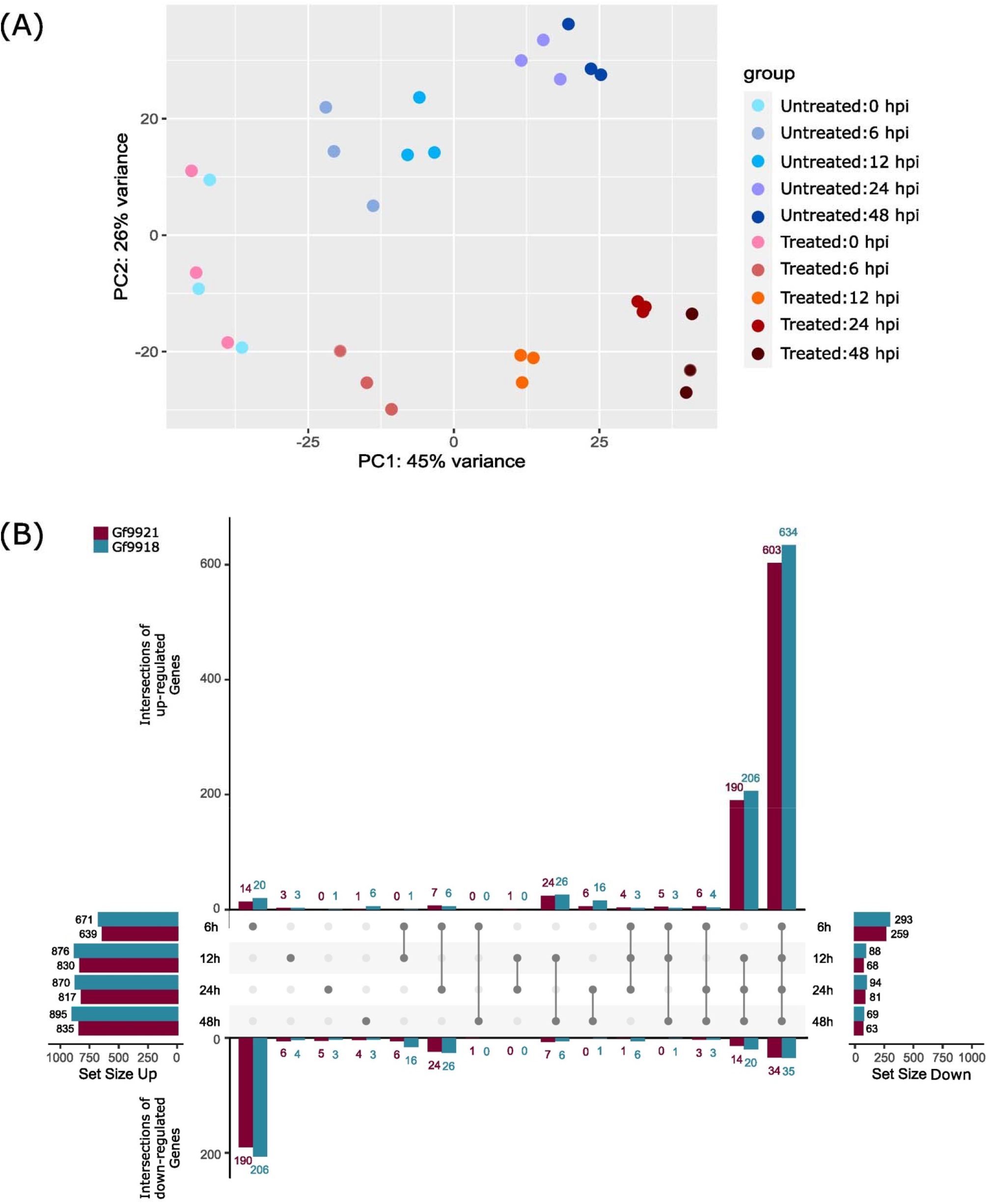
PCA plot and differentially expressed genes of the *Plasmopara viticola* infection time course experiment. **(A)** Principal Component Analysis (PCA) plot of the *P. viticola* infection experiment. Untreated samples (blue colored dots) are control samples that were not inoculated with *P. viticola* zoospores, but they were taken at the given time point. Treated samples (red colored dots) represent the inoculated samples. **(B)** Differentially expressed genes across time points. The upset plot (Lex et al., 2014; Conway et al., 2017) shows the intersection of DEGs between the time points for the up-regulated DEGs (upper part) and down-regulated DEGs (lower part) per haplotype. Data of Gf9921 are colored in magenta and data of Gf9918 in turquoise. The most left bar plot displays the total amount of up-regulated DEGs per time point and per haplotype and the most right bar plot the total amount of down-regulated DEGs per time point and per haplotype. Intersections are displayed as connecting lines with dots. The values on the y-axis represent the amount of shared DEGs.

A total of 600 to 800 up-regulated differentially expressed genes (DEGs) and 60 to 290 down-regulated DEGs per time point and per haplotype were identified using the likelihood ratio test (LRT, Figure 4 (B)). ∼600 up-regulated DEGs were found in the intersection of all time points and ∼200 up-regulated genes were shared between 12 hpi, 24 hpi and 48 hpi. Time point 6 hpi contained the highest number of down-regulated genes with ∼200.

To investigate the defense response, DEGs were functionally annotated with KAAS (https://www.genome.jp/kegg/kaas/) and the KEGG pathway ko04626 ‘Plant-pathogen interaction’ (https://www.genome.jp/dbget-bin/www_bget?ko04626) was analyzed (Figure S5). About 14 proteins of the pathway were hit by ∼31 up-regulated DEGs per haplotype and around five proteins of the pathway were hit by six down-regulated DEGs per haplotype (Table S10). Six down-regulated DEGs (see Table S10) were also found in the set of up-regulated genes. Among the up-regulated DEGs were genes encoding proteins associated with (I) production of ROS, (II) implementation of HR of pathogen associated molecular pattern (PAMP)-triggered immunity (PTI), (III) defense-related PR1 and FRK1 induction, (IV) induction of programmed cell death and (V) implementation of HR of effector-triggered immunity (ETI, Figure S5). The LRR receptor-like serine/threonine-protein kinase elongation factor Tu (EF-Tu) receptor (*EFR, K13428*)-like gene (*Gf9918_11g18310*) and the resistance to *Pseudomonas syringae* protein 2 (*RPS2, K13459*)-like genes (*Gf9918_12g19808*, *Gf9918_12g19817*) were uniquely induced in the *Rpv12*-carrying haplotype Gf9918.

### 2.7 Design of Rpv12-specific molecular markers

The new assembly with resolved *Rpv12*-alleles was exploited for marker design based on four SSR- type repeats localized within the RGA-region. Four primer pairs to amplify markers named GF14-61 to GF14-64 (Table S11) were designed and tested on different genotypes. Every marker primer pair produced a product, but only GF14-63 produced an *Rpv12*-specific allele size of 398 bp. Susceptible genotypes showed a size difference up to 448 bp (Table S12).

### 2.8 *Rpv12* locus sequence and investigation of *Rpv12* candidate genes

The *Rpv12* QTL region, delimited by SSR markers UDV-014 and UDV-370 (Venuti et al., 2013) on chr14, comprised 1.83 Mbp and 75 genes for Gf9921, 1.86 Mbp and 94 genes for Gf9918 and 2.07 Mbp and 146 genes in the corresponding genomic region of PN40024 (Figure 5).

**Figure 5.**
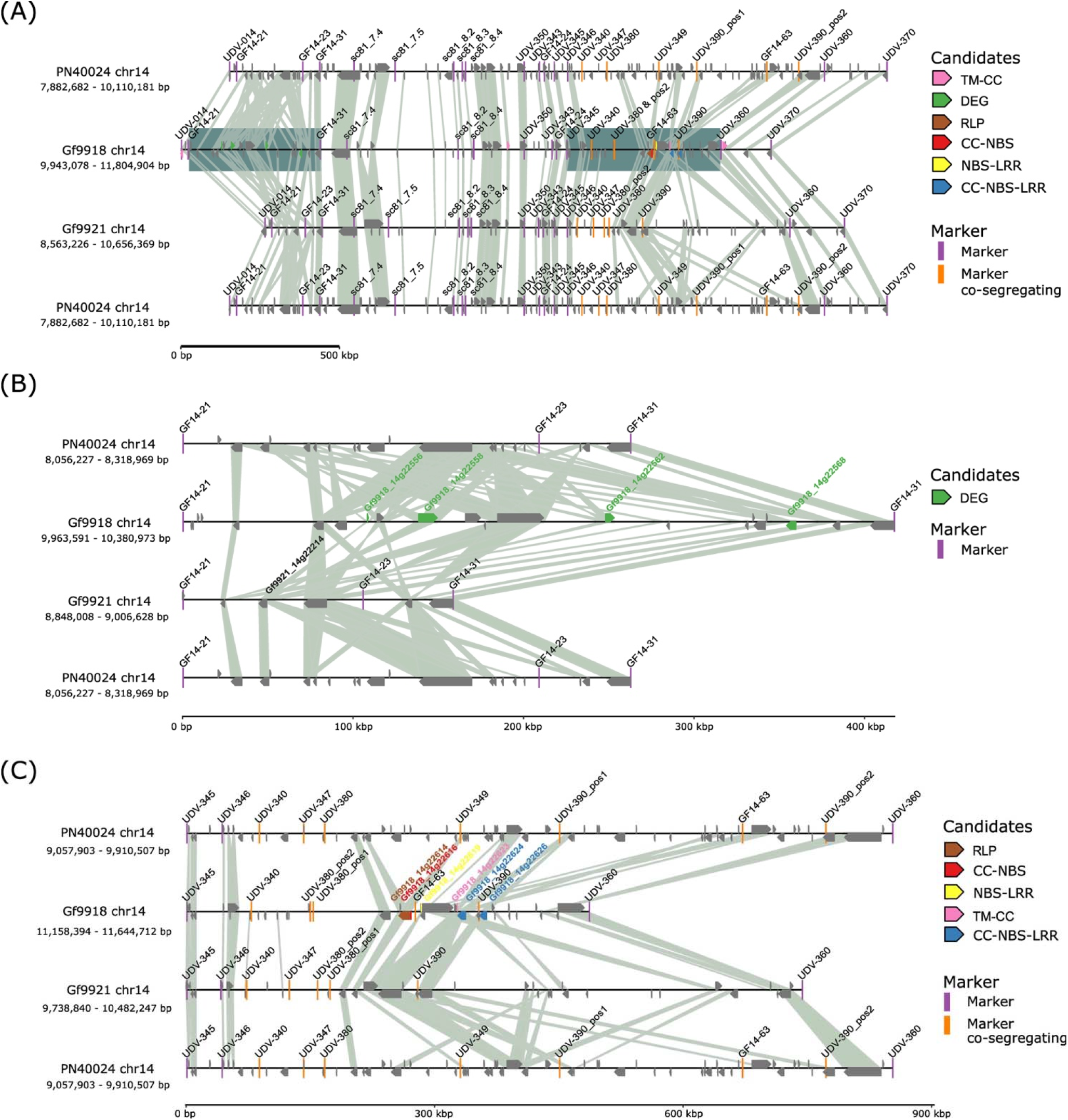
Synteny plot of the *Rpv12* locus. Connecting lines are based on blastp protein hits with a query coverage and identity >85 % and an E-value of 0.0001. To facilitate comparison, the *Rpv12* locus of PN40024 is displayed twice. **(A)** Synteny plot of the *Rpv12* locus between markers UDV- 014 and UDV-370. The two regions with candidate genes were underlain with a blue square. **(B)** Zoom in of the northern gene candidate region comprising functional candidates (markers GF14-21 to GF14-31). The gene mate *Gf9921_14g22214* for the *ACD6*-like DEGs is displayed with its name. **(C)** Zoom in of the southern gene candidate region with positional RGA candidates (markers UDV- 345 to UDV-360).

Potential *Rpv12* candidate genes were chosen from the Gf9918 gene models based on the classification as RGA by the presence of encoded resistance domains (positional candidates), or based on clear differential gene expression between susceptible and resistant haplotypes in the infection time course analysis (functional candidates). As a result, nine RGAs and five DEGs were considered as “narrow” candidates for the causal genes within the *Rpv12* locus. The candidates were analyzed for gene mates with similar protein sequences in the *Rpv12* lacking haplotype Gf9921 and the *P. viticola* susceptible genotype PN40024. Due to this restriction, the DEG *Gf9918_14g22602*, an ethylene-responsive transcription factor, was removed from the list of candidates (only two amino acid (aa) exchanges with Vitvi14g00564.t01 and Gf9921_14g22247), reducing the number of candidate DEGs to four (Table 2).

**Table 2.**
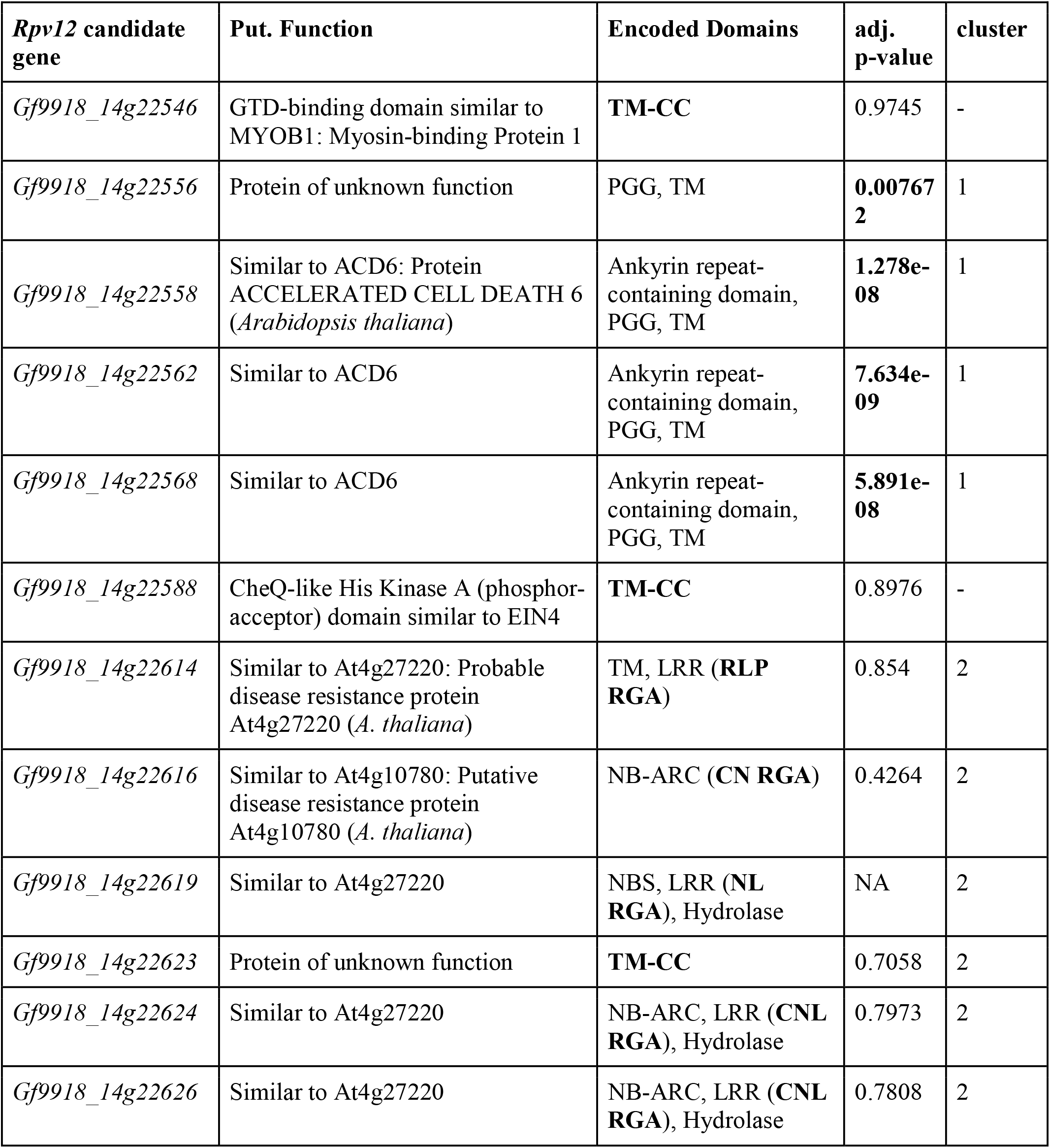
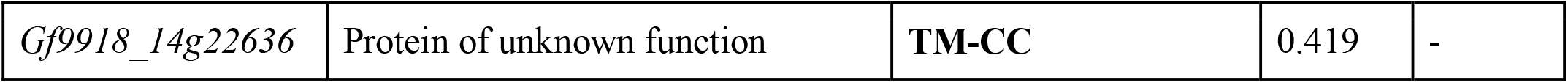
Rpv12 locus specific candidate genes of Gf9918. The designation of the Rpv12 candidate gene, the putative function (put. function) based on prediction analysis as described (gene prediction), the detected encoded domains as well as the adjusted p-value (adj. p-value) and the cluster of the locus are described. Significant adjusted p-values (p-value<0.05) and RGA classes are written in bold. For domains, the abbreviations mean the following: NBS= Nucleotide Binding Site; CNL= coiled-coil (CC), NBS and leucine-rich repeat (LRR); CN= CC and NBS; NL= NBS and LRR; RLP= Receptor like protein; TM-CC= Transmembrane and CC.

| ***Rpv12* candidate gene** | **Put. Function** | **Encoded Domains** | **adj. p-value** | **cluster** |
| --- | --- | --- | --- | --- |
| *Gf9918_14g22546* | GTD-binding domain similar to MYOB1: Myosin-binding Protein 1 | **TM-CC** | 0.9745 | - |
| *Gf9918_14g22556* | Protein of unknown function | PGG, TM | **0.007672** | 1 |
| *Gf9918_14g22558* | Similar to ACD6: Protein ACCELERATED CELL DEATH 6 (*Arabidopsis thaliana*) | Ankyrin repeat-containing domain, PGG, TM | **1.278e-08** | 1 |
| *Gf9918_14g22562* | Similar to ACD6 | Ankyrin repeat-containing domain, PGG, TM | **7.634e-09** | 1 |
| *Gf9918_14g22568* | Similar to ACD6 | Ankyrin repeat-containing domain, PGG, TM | **5.891e-08** | 1 |
| *Gf9918_14g22588* | CheQ-like His Kinase A (phosphor-acceptor) domain similar to EIN4 | **TM-CC** | 0.8976 | - |
| *Gf9918_14g22614* | Similar to At4g27220: Probable disease resistance protein At4g27220 (*A. thaliana*) | TM, LRR (**RLP RGA**) | 0.854 | 2 |
| *Gf9918_14g22616* | Similar to At4g10780: Putative disease resistance protein At4g10780 (*A. thaliana*) | NB-ARC (**CN RGA**) | 0.4264 | 2 |
| *Gf9918_14g22619* | Similar to At4g27220 | NBS, LRR (**NL RGA**), Hydrolase | NA | 2 |
| *Gf9918_14g22623* | Protein of unknown function | **TM-CC** | 0.7058 | 2 |
| *Gf9918_14g22624* | Similar to At4g27220 | NB-ARC, LRR (**CNL RGA**), Hydrolase | 0.7973 | 2 |
| *Gf9918_14g22626* | Similar to At4g27220 | NB-ARC, LRR (**CNL RGA**), Hydrolase | 0.7808 | 2 |
| *Gf9918_14g22636* | Protein of unknown function | **TM-CC** | 0.419 | - |

The four DEGs were localized at the northern end of the *Rpv12* locus in a region that is considerably larger in Gf9918 (417,382 bp) as compared to Gf9921 (158,620 bp) and PN40024 (262,742 bp) (Figure 5 (B)). This region with functional candidates differs significantly among the allelic haplotypes, yet at its borders it is flanked by highly syntenic genes.

The DEG *Gf9918_14g22556* is annotated as protein of unknown function (PUF), but a BLASTP search against the non-redundant database gave the best hit with an ankyrin repeat-containing protein ITN1-like from *V. riparia* (RefSeq XP_034707067.1) with a query coverage of 84 %, an identity of 95 % and an E-value of 4e-55. The DEG lacks the actual ankyrin repeat-containing domain and is shorter than the *V. riparia* protein (115 aa vs. 345 aa). Its expression was raised from e. g. 7.63 log2 fold change (LFC) at 6 hpi to 8.21 LFC after *P. viticola* inoculation (Figure S7 (A)).

The three DEG candidates *Gf9918_14g22558*, *Gf9918_14g22562* and *Gf9918_14g22568* were annotated as ‘Similar to ACD6: Protein ACCELERATED CELL DEATH 6’ and have a similar protein sequence and domain composition (Table 2). Even if they share almost the same protein sequence with only a few aa substitutions, they differ in their lengths due to extended start sequences of Gf9918_14g22558 and Gf9918_14g22568. Gf9918_14g22558 has a length of 715 aa, Gf9918_14g22562 of 505 aa and Gf9918_14g22568 of 692 aa (Figure S7 (A)). The three *ACD6* DEGs are positioned in around 100 kbp distance to each other. All three *ACD6* genes showed a similar expression time-course profile and had *Gf9921_22214* as gene mate (Figure S7 (B-D)). However, *Gf9921_14g22214* is allelic to *Gf9918_14g22562* with the same length of 505 aa, but with some aa substitutions.

Six of the nine positional candidate RGAs were located at the southern end of the *Rpv12* locus in a region with structural variation similar to the region with functional candidates. It contains co-segregating markers (Figure 5 (C)). Yet, different to the functional candidate region, the positional candidate region (delimited by the markers UDV-345 and UDV-360, see Figure 5) is significantly smaller in Gf9918 (486,318 bp) compared to Gf9921 (743,407 bp) and to PN40024 (852,604 bp). This region is compressed around the closely located RGAs and also flanked by highly syntenic regions.

The RGAs of the class RLP, CN, NL (NBS-LRR) and CNL shared similarity to either the *A. thaliana* disease resistance protein At4g27220 or At4g10780 (Table 2). Besides the specific RGA domains, the two CNL RGAs Gf9918_14g22624 and Gf9918_14g22626 encode two PLN03210 domains consisting of LRR motifs (Lu et al., 2020), but the domain positions, their length and the aa composition differ (e. g. Figure S8 (B)). The coding frame of Gf9918_14g22626 is lacking 113 aa followed by another lack of 42 aa within a LRR domain in comparison to Gf9918_14g22624.

Additionally, the *Rpv12* locus contained four TM-CC (Transmembrane and Coiled-Coil) RGAs. Two TM-CCs, *Gf9918_14g22623* inside of the candidate region with positional candidates and *Gf9918_14g22636* southern of this region, were annotated as ‘protein of unknown function’. The other TM-CCs were functionally annotated as containing a ‘globular-tail (GTD)-binding domain similar to Myosin-binding Protein 1 (MYOB1)’ (*Gf9918_14g22546*, *Rpv12* start) and as containing a ‘CheQ-like His Kinase A (HisKA) domain similar to ETHYLENE INSENSITIVE 4 (EIN4)’ (*Gf9918_14g22588*, *Rpv12* mid).

Considering the expression, significant high counts were found for the CNLs and the TM-CC *Gf9918_14g22588* under similar expression of treated and untreated samples. In contrast, the TM-CC *Gf9918_14g22636* had ∼300 normalized counts and the RLP had up to 100 counts. The other RGAs had no or only a few counts. None of the RGAs was differentially expressed.

## 3 Discussion

### 3.1 The pedigree of *Rpv12* carrier Gf.99-03

The goal of this work was to characterize the grapevine *Rpv12* locus on chr14 which confers resistance to the downy mildew pathogen *P. viticola*. Since the *Rpv12* locus displays dominance and is often present in heterozygous state, a phase-separated genome sequence was required. The separation of both haplotypes was possible by using the tool TrioCanu (Koren et al., 2018), but necessitates sequence information of both parental genotypes of the targeted heterozygous F1. Thus, access to a complete ‘trio’ with correct descendance is necessary as given for genotype Gf.99-03 that was selected for analysis.

The pedigree of Gf.99-03 was confirmed, including parentage of the genotypes Gf.43-21 and 65-153- 18 as well as some contributions of the grapevine varieties ‘Blaufraenkisch’, ‘Calandró, ‘Regent’ and ‘Dominá as known from breeding records. No relationship was demonstrated between the genotype Gf.99-03 and the cultivars ‘Kunbarat’ or ‘Kunleaný, an expected result due to genetic distance. Based on Koleda *et al*. (Koleda, 1975), the parental genotypes of ‘Kunbarat’ and 65-153-18 originate from the same pedigree. Regrettably, the interspecific genotypes 28/19 and 4/15, of which 4/15 is expected to be one parent of 65-153-18 (Figure 1), are not available anymore prohibiting analysis.

All studied genotypes were checked for the presence of further known and marker-tagged resistance loci in addition to *Rpv12*. Based on SSR marker analysis, no other characterized *Rpv* locus was detected in Gf.99-03.

### 3.2 Gf.99-03 is a highly *P. viticola* resistant *Rpv12-*carrier

The effectiveness of *Rpv12* associated resistance properties was previously demonstrated (Wingerter et al., 2021). Gf.99-03 responded to *P. viticola* challenge by production of ROS within 24 hpi. The *Rpv12-*carrying genotype 65-153-18 allowed less than half of the mycelial growth observed in susceptible ‘Italiá controls. The inhibition of mycelial development was even more pronounced in its descendant Gf.99-03 (*Rpv12*). Nevertheless, repression of mycelial expansion of *P. viticola* was strongest in the maternal genotype Gf.43-21 that carries two different *P. viticola* resistance loci, namely *Rpv1* and *Rpv3.1*. *Rpv1* is a strong resistance locus from *Muscadinia rotundifolia* (Merdinoglu et al., 2003), and *Rpv3.1* (Welter et al., 2007) is an American *Vitis* spec. locus most likely derived from *V. rupestris* (Rockel et al., 2021). However, marker analyses confirmed that neither *Rpv1* nor *Rpv3.1* were inherited to Gf.99-03.

Although Gf.99-03 carries only *Rpv12* as known resistance locus against *P. viticola*, the effect of this QTL on mycelial growth seems to be more inhibiting in Gf.99-03 than in its parent 65-153-18. It is possible that unrecognized minor resistance factors were inherited to Gf.99-03, which contribute to its stronger defense of *P. viticola*. Some other weak resistance QTLs to *P. viticola* were described in *V. amurensis* ‘Shuang Honǵ, but remain as yet uncharacterized (Fu et al., 2020).

The genotype Gf.99-03, besides carrying the locus *Rpv12*, contains resistance QTLs directed against powdery mildew (*Erysiphe necator*, *Ren3* and *Ren9*) on chr15 (Welter et al., 2007; Zendler et al., 2017; Zendler et al., 2021). It is possible that defense genes within different resistance loci interact and reinforce each other. It is known from the *M. rotundifolia* resistance QTL *Run1/Rpv1* that genes in close proximity act against *P. viticola* and *E. necator* (Feechan et al., 2013).

Overall, the *Rpv12-*carrier Gf.99-03 was validated as a highly *P. viticola* resistant genotype. However, it allowed the formation of haustoria in the early phase of pathogen attack (72 hpi), indicating post-invasion and post-haustorial resistance. The *P. viticola* haustoria on Gf.99-03 were more numerous and much smaller as compared to a susceptible genotype, appearing rudimentary or atrophied (Figure 2 (G, H)). The formation of haustoria plays a central role in the infection process of *P. viticola*. It was shown that haustoria formation is stopped in non-host plants and that defense reactions are triggered in resistant grapevine varieties as soon as haustoria are detected (Diez-Navajas et al., 2008). Formation of haustoria may therefore be a prerequisite to initiate defense reactions in the resistant host. In *A. thaliana* the TM-CC proteins RPW8.1 and RPW8.2 mediate broad spectrum resistance to the powdery mildew pathogen *Golovinomyces* spp. (Xiao et al., 2001; Wang et al., 2009; Kim et al., 2014). The RPW8.2 protein is transferred to the extrahaustorial membrane and activates a salicylic acid-dependent defense pathway (Xiao et al., 2003). It leads to ROS accumulation in the vicinity of haustoria presumably to constrain their development (Wang et al., 2009). ROS production was also observed here during *Rpv12* mediated defense, as also in an earlier investigation on the *Rpv3* locus from an American *Vitis* species (Eisenmann et al., 2019; Rockel et al., 2021). A previous study on *P. viticola* defense reaction of the *V. riparia* accession ‘Gloire de Montpellier’ identified an induced gene family of three *VRP1* CNL genes *VRP1-1, 1-2 and 1-3* located on chr10 (Kortekamp et al., 2008), apart from its resistance QTLs *Rpv5* on chr9 and *Rpv6* on chr12 (Marguerit et al., 2009). Astonishingly, the proteins encoded by *VRP1-1* and *VRP1-2* include an N-terminal domain with similarity to RPW8 (Kortekamp et al., 2008), opening the hypothesis that similar reaction mechanisms may be active in grapevine and in *A. thaliana* during defense against different mildews.

### 3.3 Phased genome assembly and gene annotation

The Gf.99-03 diploid genome assembly represents one of the few high-quality and truly phase-separated *Vitis* genome sequence assemblies currently available. The overall assignment to pseudochromosomes was validated *in silico* with SSR markers (Table S1). The Gf9918 haplotype assembly contains around 33 Mbp more than the Gf9921 haplotype. The additional sequences are related to different dicotyledon species and close relatives of *Vitis* (clade Eudicots, clade Rosids or order *Vitales*). Already Lodhi and Reisch (Lodhi and Reisch, 1995) reported varying genome sizes of 439 to 526 Mbp for Asian *Vitis* species and 411 to 541 Mbp for North American *Vitis* species.

Considering the reported *Vitis* genome sizes and the descent of Gf.99-03’s parent 65-153-18 from Asian *V. amurensis*, the increased genome size of the paternal haplotype Gf9918 is not unexpected. Despite the higher base content of Gf9918, the gene annotation and RGA annotation of the haplotype assemblies resulted in an even set of gene models and RGAs.

### 3.4 Differentially expressed genes

The analysis of DEGs was based on an infection time-course experiment using *P. viticola* infected leaf disc samples. The number of sequenced-tagged mRNAs across the various infection stages was approximately equal and the number of genes with a count was similar in both haplotypes. PCA analysis indicated a successful infection time-course experiment with the data points well separated according to condition, time and treatment. As expected, the treated samples of time point 0 hpi when the *P. viticola* infection was initiated, but had no time to spread, clustered closely with the data points of untreated time point 0 hpi. The differential gene expression analysis resulted in a higher amount of up-regulated DEGs than down-regulated DEGs and equal amounts of DEGs for each haplotype and per time point. It is noticeable that the number of low expressed DEGs was around 250 (Gf9921) to 300 (Gf9918) at time point 6 hpi and decreased to a stable number of 60 to 90 DEGs for the other time points.

During the defense response of Gf.99-03 against *P. viticola*, ∼31 genes were up-regulated whose proteins putatively play a role in plant-pathogen interactions, namely in ROS production, during HR induced by PTI or ETI, and in other defense-related pathways. Genes similar to *AtRPS2* (on chr12) and *EFR* (on chr11) were exclusively induced in the *Rpv12*-carrying haplotype upon *P. viticola* challenge.

*AtRPS2* genes have been described as plant disease resistance genes carrying LRR domains (Bent et al., 1994) and more recently as NLR motif encoding gene, which interacts with bacterial effector protein AvrRpt2. Overexpression of *AtRPS2* showed higher accumulation of ROS (Li et al., 2019). In *Oryza sativa*, it confers resistance to fungal and bacterial pathogens like *Magnaporthe oryzae* and *Xanthomonas oryzae* (Qi et al., 2011; Li et al., 2019). In this study, the transcript-isoforms of the *RPS2*-like gene *Gf9918_12g19808* carry either all protein domains for NLR or only LRR domains.

*EFR* genes (receptor kinases), showing motifs of serine-threonine protein kinases, are described as interacting with bacterial EF-Tu and activating the basal immunity (Zipfel et al., 2006). Here, it may be involved in the interaction with an oomycete.

The identification of up-regulated genes involved in PTI and ETI supports the plant-oomycete interaction model described in Judelson and Ah-Fong (2019), where oomycete infection is detected through pattern recognition receptors (PRRs) in the intercellular mesophyll space (Judelson and Ah-Fong, 2019). The up-regulated genes of the infection time-course experiment indicate a complex oomycete detection causing a multi-level defense response. As *Rpv12* describes ∼79 % of the resistance phenotype (Venuti et al., 2013), the *Rpv12* locus must contain specific genes that significantly contribute to the *P. viticola* resistance.

### 3.5 *Rpv12* locus and potentially resistance related genes

The sequence of the *Rpv12* locus was extracted from both haplotypes of Gf.99-03, namely from the *Rpv12* negative haplotype sequence Gf9921 and the *Rpv12-*carrying haplotype sequence Gf9918. Comparison of these sequences and their gene content was extended to the homologous region of the susceptible reference model grapevine genotype PN40024 (Jaillon et al., 2007; Canaguier et al., 2017), revealing considerable structural differences and partial hemizygosity. To identify *Rpv12* associated genes, RGAs encoding typical protein domains as well as DEGs during pathogen challenge in the locus were studied.

Resistance research in *Vitis* has so far been focused on CNL/TNL genes, especially in the context of the *Rpv12* locus (Di Gaspero and Cipriani, 2003; Venuti et al., 2013; Chitarrini et al., 2020). CNL and TNL genes represent the largest class of *R*-genes known in the plant kingdom (Dangl and Jones, 2001; Hulbert et al., 2001; Wei et al., 2020) and they play an important role in pathogen defense. To follow the recent plant-oomycete interaction model (Judelson and Ah-Fong, 2019), the focus was on the PRRs, RLPs and RLKs monitoring the extracellular space and the TM-CCs. The complete region between the QTL delimiting SSR markers UDV-014 and UDV-370 was analyzed (Venuti et al., 2013). R-Genes are often organized in clusters (Hulbert et al., 2001; Meyers et al., 2003). Two clusters of typically resistance-associated genes were identified in this region. These clusters are structurally highly diverse between the *Rpv12* carrying haplotype Gf9918 and the *Rpv12* non-carrying haplotype Gf9921 as well as compared to the susceptible PN40024 genotype. They differ in sequence size and presence of resistance-associated genes present in Gf9918, but are flanked by highly syntenic gene regions. One cluster, significantly larger in Gf9918 (Figure 5 (B)), includes three copies of *ACD6*-like R-genes that show up as DEGs and encode Ankyrin repeat-containing domains, PGG and TM domains. The three *ACD6*-like genes *Gf9918_14g22558*, *Gf9918_14g22568* and *Gf9918_14g22562* were found within a region of ∼221 kbp. In *A. thaliana*, a hyperactive allele of *AtACD6* (*At4g14400*) conferred resistance to various pathogens including microbes and insects (Todesco et al., 2010), making these genes important candidates for *P. viticola* resistance in grapevine. Also, a DEG (*Gf9918_14g22556*) encoding a protein of unknown function containing a PGG and a TM domain was identified.

The locus was further described by co-segregating markers between marker UDV-350 and UDV-370 (Venuti et al., 2013) reducing the size to 1.01 Mbp and 46 genes in Gf9921, to 0.78 Mbp and 51 genes in Gf9918 and to 1.14 Mbp and 88 genes in PN40024. Here, positional RGA candidates were found. The co-segregating marker GF14-63, that was designed in this study for downstream marker assisted breeding application, was placed between the first three RGAs. The marker UDV-390 is positioned between the other three RGAs.

The RLP gene *Gf9918_14g22614*, carrying TM and LRR domains and similar to the NL Gene *Gf9918_14g22619*, is a homolog of the *A. thaliana* gene *At4g27220. At4g27220* is orthologous to *GbaNA1* that was described as a resistance mediating gene of *Gossypium barbadense* against *Verticillium dahlia*. It activates ROS production and ethylene signaling in *A. thaliana* (Li et al., 2018). Besides, *Gf9918_14g22624* and *Gf9918_14g22626*, encoding for proteins with CC, NB-ARC and LRR domains (CNL), are orthologous to *At4g27220*. Additionally, they carry PLN03210 domains which were originally found in proteins of the TNL class like RPS6 or RAC1 (Kim et al., 2009). RAC1 mediates resistance to the oomycete *Albugo candida* in *A. thaliana* (Borhan et al., 2004; Kim et al., 2009). The orthologue *At4g10780* of the CN gene *Gf9918_14g22616* carries an RPS6 domain, while *Gf9918_14g22616* itself only carries an NB-ARC domain. For the TM-CC gene *Gf9918_14g22623,* directly upstream of the CNL gene *Gf9918_14g22624,* no similarities were found.

The gene *Gf9918_14g22588*, encoding a TM-CC protein, is located in the region between the two clusters and putatively codes for a CheQ-like histidine kinase A (phosphate-acceptor) domain similar to EIN4. AtEIN4 (At3g04580) is known as an ethylene receptor negatively regulating ethylene signaling (Hua and Meyerowitz, 1998) that also displays a serine/threonine kinase activity *in vitro* (Moussatche and Klee, 2004). It is located on the membrane of the endoplasmic reticulum (ER) and the histidine kinase domain is active in signal transduction. Thus, the TM-CC protein encoded by *Gf9918_14g22588* may function in defense-related ethylene signaling.

In conclusion, the *P. viticola* resistance locus *Rpv12* of grapevine mediates post-invasion, post haustorial resistance. The pathogen may form multiple but atrophied haustoria on resistant plants, likely due to the struggle of the oomycete to gain access to plant nutrients, but inhibited by defense reactions of the host cells including the formation of ROS. The defense reaction leads to localized necrosis and HR. This observation agrees with the clearly pathogen-induced expression of three *ACD6*-like genes present in one cluster of the *Rpv12* locus. A second cluster contains several RGAs, whereof *Gf9918_14g22624* and *Gf9918_14g22626* are constitutively expressed. These genes encode CNL domain proteins and may act as primary receptors of pathogen-derived signals (PAMPs) to recognize the attack by *P. viticola*. In addition, further resistance-associated gene functions were identified in the *Rpv12* region. Considering the whole genomic region, the resistant haplotype shows extensive structural divergence when compared to the susceptible haplotype and to the genome sequence of PN40024, the susceptible *V. vinifera* model genotype.

This situation is reminiscent of a recent local haplotype resolution and characterization of the *M. rotundifolia* ‘Trayshed’ QTLs *Run1.2* and *Run2.2* directed against *E. necator* (powdery mildew), which revealed quite substantial structural variation of potential disease resistance genes in the QTL regions (Massonnet et al., 2022). Despite this divergence of genomic regions carrying resistance loci in wild American or Asian *Vitis* relatives (or the closely related American *M. rotundifolia*), there seems to remain enough synteny to the noble European *Vitis vinifera* cultivars genomes to allow the introgression of QTL loci to generate naturally resistant grapevines for viticulture. Stacking of several independent loci is required to enhance sustainability of the resistance traits. To this end, tightly QTL-linked molecular tags – ideally linked to the responsible resistance genes- are required.

In this work, a new SSR marker, GF14-63, localized in the center of the polymorphic *Rpv12* region and between the two RGA clusters was developed. This marker should be highly useful in marker assisted resistance breeding of genetically resistant and more sustainable grapevine varieties. Regarding the *Rpv12* mediated resistance pathway, it can be suspected that *P. viticola* penetration is detected inside the cell by one of the two constitutively expressed CNL genes, but that the differentially expressed *ACD6*-like genes contribute to exert the local necrosis, and the resistance cascade is initiated in concert with additional regulated functions spread over the chromosomes. Other RGAs identified within the *Rpv12* locus may also play an important role.

## 4 Material and Methods

### 4.1 Plant material

Plant material of Gf.99-03 (Gf.2014-099-0003, *V*IVC <u>27131</u>), Gf.43-21 (Gf.2004-043-0021, *Rpv1*, *Rpv3.1*, *Ren3*, *Ren9, V*IVC <u>27130</u>) and 65-153-18 (*Rpv12, V*IVC <u>41129</u>) was taken from the germplasm collection of Julius Kühn-Institute, Institute for Grapevine Breeding Geilweilerhof (49°12’54.1’’N, 8°02’41.3’’E). Wooden two eye cuttings were rooted in Jiffy-7® (Jiffy Products International BV, Zwijndrecht, Netherlands) pots and propagated in the greenhouse. While genotype 65-153-18 (4/15 x ‘Blaufraenkisch’) is an offspring of an Hungarian breeding line based on *V. amurensis* (Koleda, 1975), Gf.43-21 (‘Calandro’ x VRH3082 1-49) results from a series of crosses, started in 1916 with backcrosses of *Muscadinia rotundifolia* (Olmo, 1986) (https://www.vivc.de/).

### 4.2 gDNA extraction for SSR marker analysis

Genomic DNA (gDNA) of the genotypes Gf.43-21, Gf.99-03, 65-153-18 as well as ‘Lemberge’ (*V*IVC <u>1459</u>), ‘Calandró (*V*IVC <u>21797</u>), ‘Regent’ (*V*IVC <u>4572</u>) and ‘Dominá (*V*IVC <u>3632</u>) was isolated. Small pieces (0.25 cm^2^) from the first apically inserted leaf were collected, precooled in 1.5 ml reaction tubes (Eppendorf, Hamburg, Germany), lyophilized (Martin Christ Gefriertrocknungsanlagen GmbH) and ground in a paint shaker (SK450, Fast & Fluid Management B. V., Sassenheim, Netherlands) with metal beads (3 mm in diameter, Ditzinger, Brunswick, Germany). DNA was extracted using the NucleoSpin 96 Plant Core Kit (Macherey Nagel).

### 4.3 SSR marker analysis

The sequences of primer pairs flanking SSR markers used in this work are listed in Table S1. All oligos were synthesized by Metabion International, Planegg, Germany. PCR amplification and marker measurement were done as described (Müllner et al., 2020). A marker set of 203 primer pairs was applied to gDNA of Gf.99-03, Gf.99-21 and 65-153-18.

### 4.4 *P. viticola* inoculation of leaf discs for a time course of transcriptomic analysis and follow-up by microscopy

Leaves for high molecular genomic DNA and RNA extraction were taken early in the morning to avoid inhibition of purification steps by starch accumulation.

From three clonally replicated plants of Gf.99-03 two leaves were taken from the third and fourth apical insertion and cleaned in deionized water. Leaf discs of 1.5 cm diameter were punched out by a surface sterilized cork borer. In total, forty leaf discs were tested per plant. They were placed on 1 % agar in H_2_O_dest_ (Gustav Essig GmbH & Co. KG, Mannheim) in rectangular light-transmissible incubation plates covered with a lid (Corning 431111, Corning Incorporated, Corning, NY, USA). 20 leaf discs were mock-inoculated with 40 µl H_2_O_dest_ and 20 leaf discs were inoculated with 40 µl of a *P. viticola* zoospore suspension. To obtain these zoospores, sporangia (20.000/ml) were diluted in H_2_O_dest_ and stirred every 15 min until zoospore release as controlled by bright field microscopy. Zoospores were separated from sporangia with a filtration step as described (Müllner and Zyprian, 2022). Inoculated leaf discs and mock controls were kept at 22 °C and 100 % humidity with a photoperiod of 16 h light and 8 h darkness. The ongoing infestation of all samples was checked by fluorescence microscopy. At every time point (0, 6, 12, 24, 48 hpi), four leaf discs per sample were taken, punched again by a smaller 1.3 cm diameter cork borer, shock frozen in liquid nitrogen and stored at −70 °C until RNA extraction.

### 4.5 Fluorescence microscopy to follow the infestation progress

Parallel studies on additionally inoculated leaf pieces were done to observe mycelial ingress. At 24 and 48 hpi, pieces of tested leaves were bleached in 1 N KOH for at least 2 h at 65 °C. Subsequently, samples were submitted to alkaline aniline blue staining as described (Hood and Shew, 1996; Müllner and Zyprian, 2022).

### 4.6 Detection of reactive oxygen species

ROS were revealed by Nitrotetrazolium blue chloride (Merck KGaA, Darmstadt, Germany) staining according to the protocol of (Kumar et al., 2014) and necrosis around stomata was detected with Auramine O und Calcofluor white staining. For details see Methods S1.

### 4.7 Assessment of mycelial growth

Mycelial development was followed to study the inhibition of pathogen progress mediated by the *Rpv12* locus after 72 hpi by comparison of genotypes Gf.43-21, Gf.99-03 and 65-153-18. The cultivar ‘Italiá served as susceptible control. For every genotype under study, three leaves were taken and out of every leaf three leaf discs were stained with alkaline aniline blue (see above). Of every leaf disc, five pictures of 100x magnification were taken using a fluorescence microscope with GFP-Filter (Leica A). In total, 45 images were taken at 72 hpi from every genotype and analyzed (Müllner and Zyprian, 2022). The area of mycelium was determined using Welch’s-T-Test and R Software v4.0.3 (Team, 2018) (https://www.R-project.org/).

### 4.8 RNA extraction of Gf.99-03 tissues and the leaf discs from the *P. viticola* inoculation time course experiment

RNA was extracted from leaf discs, whole leaves, tendrils, roots and stems of Gf.99-03. Samples were ground using mortar and pistil (Jipo, Czech Republic) with liquid nitrogen. To avoid potential contamination by plant phenolics, a pinch of PVP-40 (Sigma Aldrich, Merck KGaA, Darmstadt, Germany) was added to the samples. RNA was isolated using the Spectrum^TM^ Plant Total RNA Kit (Sigma Aldrich, Merck KGaA, Darmstadt, Germany). The material was stored at −70 °C until use.

### 4.9 DNA extraction for PacBio Sequencing of Gf.99-03 and for Illumina sequencing of Gf.99- 03, Gf.43-21 and 65-153-18

High molecular weight genomic DNA was extracted from 2.5 g fresh leaf samples. The material was transferred into liquid nitrogen and ground in a mortar. Genomic DNA was extracted from young leaves using the CTAB method (Rosso et al., 2003). The samples were purified using QIAGEN Genomic-tip 500/G.

### 4.10 Library construction and sequencing

The Gf.99-03 DNA samples were sequenced on a Sequel I sequencer. The PacBio Sequel libraries were prepared, and samples sequenced on two runs using sequencing chemistry 3.0, binding kit 3.0 and DNA Polymerase 3.0 on 1Mv3 SMRT cells (Pacific Biosciences). The subread BAM files were converted to FASTA format with SMRT Link v5.1.0.26412 (PacBio Reference Guide 2018) and reads smaller than 500 bp were removed.

The gDNA samples of Gf.99-03, Gf.43-21 and 65-153-18 were sequenced 2x150 bp paired-end on an Illumina NextSeq-500 (Illumina, San Diego, USA). Illumina NGS libraries were created following the TruSeq DNA Sample Preparation v2 Guide. The Illumina short read data were trimmed with Trimmomatic-v0.39 (Bolger et al., 2014) with the parameter ‘LEADING:34 TRAILING:34 SLIDINGWINDOW:4:15 ILLUMINACLIP:2:34:15 MINLEN:90’.

The mRNA tissue samples of tendrils, roots and stems and the mRNA samples of the infection time course experiment were sequenced 2x75 bp single-end on a NextSeq-500. The RNA-Seq libraries were prepared using 1,000 ng total RNA following the TruSeq RNA Sample Preparation v2 Guide. The RNA-Seq data were trimmed with Trimmomatic-v0.39 removing reads shorter than 60 bp length. The other parameters were the same as for DNA data.

An RNA sample of Gf.99-03 leaves was sent to CeGaT GmbH Tübingen. The preparation of the RNA-Seq library was done using 1,000 ng of total RNA according to TruSeq Stranded mRNA (Illumina) Kit and sequenced 2x100 bp paired-end on an Illumina NovaSeq-6000. The data were trimmed with Trimmomatic-v0.39 allowing a minimal read length of 80 bp. The other parameters were set as for the genomic short read data.

### 4.11 Phase-separated genome assembly and pseudochromosome construction

The Gf.99-03 long reads were phase separated based on their 24-mer profile with the TrioCanu module of Canu v1.9 (Koren et al., 2017; Koren et al., 2018). As input, the Illumina reads of Gf.99- 03s’ parental genotypes Gf.43-21 and 65-153-18 together with the long reads of Gf.99-03 were given to TrioCanu and their 24-mer profile computed. According to the k-mer comparison, the long reads were binned into the parental read subsets and one unassigned subset (bins). The usage of 24-mers was empirically tested by checking the average and maximal read length per parental bin and a small bin size of the unassigned reads for different k-mers. K-mer sizes varying from 18-mers to 27-mers were tested (data not shown).

To compute the haplotype assemblies of Gf.99-03, named Gf9921 (maternal haplotype) and Gf9918 (paternal haplotype), both parental read-subset were individually assembled with Canu v1.9 with the settings ‘corMhapSensitivity=normal’, ‘correctedErrorRate=0.085’ and ‘genomeSize=500m’. All steps were performed on a compute cluster. The assembly parameters were chosen through evaluating different assembly runs. The number of sequences, the total assembly length and the repeat length all reported by Canu or determined with QUAST v5.0.2 (Mikheenko et al., 2018) as well as the plant core genes predicted with the program BUSCO v3.0.2 utilizing the pre-release of the database ‘eudicots_odb10’ comprising 2,326 plant core genes (Waterhouse et al., 2018) were considered. The parameters resulting in haplotype assemblies having the smallest distance in these values were selected. The Gf9921 and Gf9918 haplotype assemblies were polished two times with *arrow* (gcpp 1.0.0-SL-release-8.0.0; smrtlink_8.0.0.80529) using the haplotype-specific read bin. Around 25 contigs <10 kbp were discarded.

Polished contigs were assigned to pseudochromosomes based on RBHs with the PN40024 12X.v2 (Jaillon et al., 2007; Canaguier et al., 2017), the *V. riparia* Gloire de Montpellier genome assembly (Girollet et al., 2019) and with the ‘Börner’ haplotype assemblies (Frommer et al., 2022). Therefore, a gene prediction was performed on Gf9921 and Gf9918 with AUGUSTUS v3.3 (Stanke and Waack, 2003) using the generated PN40024 parameter file from (Frommer et al., 2022), ‘--UTR=on’ and ‘-- allow_hinted_splicesites=atac’. The protein sequences of these gene predictions were used in a BLASTP search against the longest protein sequences per gene of the PN40024 VCost.v3 gene annotation and against the *V. riparia* and ‘Börner’ longest protein sequences and *vice versa*.

Additionally, a BLASTN search of the contigs against the pseudochromosomes of these three genotypes and *vice versa* was performed. RBHs were constructed from the blast hits with maximal E- value of 0.0001 for both directions and ≥ 80 % coverage and ≥ 80 % identity for at least one direction. A contig was only assigned through protein RBHs if it had more than 10 RBHs with a pseudochromosome. If two or more pseudochromosomes had RBHs with a contig, the pseudochromosome with the most RBHs was chosen if the RBH amount was higher than 30 %.

For the formation of pseudochromosomes, protein RBHs with PN40024 were given higher preference than the protein RBHs with the other two cultivars and finally the nucleotide RBHs. Contradicting assignments to different pseudochromosomes of the cultivars were solved through choosing the pseudochromosome of the cultivar showing the most protein RBHs.

Based on the position of the RBHs on the pseudochromosome, the contigs were arranged into pseudochromosomes and connected with 100 N’s. To further assign remaining contigs, reciprocal BLASTN hits were iteratively computed between the contigs and the pseudochromosomes of the other haplotype (e. g. Gf9921 contigs against Gf9918 pseudochromosomes). These construction and assignment steps were repeated until no more contigs could be assigned.

To verify the orientation and order of the contigs on the pseudochromosomes, dot plots of 1-to-1 alignments between the homologous pseudochromosomes of the Gf.99-03 haplotypes and dot plots between the haplotypes and the PN40024 12X.v2 pseudochromosomes were generated with DNAdiff v1.3 and mummerplot v3.5 of the MUMmer package v4.0.0beta2 (Marcais et al., 2018). 470 SSR- repeats (Table S1) were identified on the Gf.99-03 and PN0024 genome sequence assemblies with primer search of the EMBOSS v6.6.0.0 package (Rice et al., 2000) allowing 20 % mismatches. These bioinformatically revealed marker positions were depicted on the dot plots. Contig position, orientation and the pseudochromosome assignment was adapted if the dot plots and SSR markers showed a disagreement between the haplotypes and between the haplotypes and PN40024. Furthermore, contigs on which markers of two different chromosomes mapped, were investigated for mis-assemblies. All-versus-all dot plots generated with D-Genies v1.2.0 (Cabanettes and Klopp, 2018) using minimap2 (Li, 2018) and mappings of the haplotype-specific genomic reads computed with minimap2 v2.17 (‘-ax map-pb --secondary=no’) were examined for breakpoints. If a sequence region within a contig was found where the read coverage was dropping to less than five and if the all-versus-all dot plot and the SSR markers agreed and would assign the parts of the contig sequence to another pseudochromosome, the contig was split and the sequences assigned according to the data, see Methods S2.

A BUSCO analysis was computed on the Gf.99-03 haplotype assemblies at pseudochromosome level with BUSCO v5.1.2 and the ‘eudicots_odb10’ dataset (Seppey et al., 2019). The pseudochromosome lengths were visualized with cvit v1.2.1 (Cannon and Cannon, 2011). To validate the genetic purity of the Gf.99-03 assembly, the Gf.99-03 long reads as well as the Gf9921 and Gf9918 haplotype assembly were investigated for contaminations through a BLASTN search against the NCBI nucleotide database v4 (parameter ‘-evalue 0.0001 -max_target_seqs 1 -max_hsps 1’).

### 4.12 Examination of haplotype-phasing

19-mer databases were computed for the Illumina read data of the parent-child trio (Gf.99-21, 65- 153-18, Gf.99-03) with Merqury v1.3 (Rhie et al., 2020) and haplotype specific k-mer sets (hapmers) were identified. The contig sequences of the haplotype assemblies were evaluated setting the parameters ‘num_switch 10’ and ‘short_range 20,000’.

### 4.13 Gene prediction

For both Gf.99-03 haplotypes, an individual repeat library was generated according to the MAKER ‘Advanced Repeat Library Construction Protocol’ (Campbell et al., 2014) using the settings mentioned in the protocol.

Additionally, a *de novo* and genome-guided transcriptome assembly were computed from the paired-end RNA-Seq leaf data with Trinity v2.10.0 (Grabherr et al., 2011). First, the trimmed RNA-Seq data were mapped with GMAP/GSNAP version 2020-09-12 (Wu and Watanabe, 2005) on Gf9921 and on Gf9918 using ‘-B 5 --novelsplicing 1’. The trimmed fastq files were used for the Gf.99-03 *de novo* transcriptome assembly and the primary mappings were processed into a Gf9921 and Gf9918 genome-guided transcriptome assembly using the parameter ‘-- genome_guided_max_intron 20000’.

*Ab initio* gene predictions were carried out on each haplotype with GeneID v1.4.5-master- 20200916 with the publicly available *V. vinifera* parameter set (vvinifera.param.Jan_12_2007), GlimmerHMM v3.0.4 (Majoros et al., 2004), SNAP v2006-07-28 (Korf, 2004) and BRAKER2 v2.1.5-master_20201122 (Hoff et al., 2016; Hoff et al., 2019; Bruna et al., 2021). For GlimmerHMM training, a gene prediction training set was generated with PASA v2.4.1 (Haas et al., 2003) and TransDecoder v5.5.0 (https://github.com/TransDecoder/TransDecoder) according to the PASA wiki (https://github.com/PASApipeline/PASApipeline/wiki/). For the generation of PASA alignment assemblies, the Trinity *de novo* and genome-guided transcriptome assemblies, *Vitis* full-length cDNA sequences (NCBI, Nucleotide DB “Vitis” [Organism] AND complete cds[Title], downloaded on 05.06.2020) and *Vitis* EST sequences (NCBI, txid3603[Organism:exp] AND is_est[filter], downloaded on 27.07.2020) were used. The parameters for PASA were the same as in (Frommer et al., 2022). GlimmerHMM was trained with 7,500 (Gf9921) and 5,500 (Gf9918) PASA/TransDecoder genes and a GlimmerHMM gene prediction performed.

SNAP training was performed three times with MAKER v3.01.03 (Holt and Yandell, 2011; Campbell et al., 2014). First, the Trinity *de novo* and genome-guided transcriptome assemblies, the *Vitis* full-length cDNA sequences, *Vitis* protein sequences (NCBI, Protein DB “Vitis”[Organism], downloaded on 05.06.2020), Viridiplantae sequences (UniProt, release 2020_02, “Viridiplantae [33090]” AND reviewed:yes), RepBase monocotyledons repeat library (RepeatMaskerEdition- 20181026) (Bao et al., 2015), the generated haplotype specific repeat library and MAKER transposable element sequences were aligned and processed with MAKER using Exonerate v2.4.0 (Slater and Birney, 2005) and genes derived with MAKERs internal algorithm using the parameter ‘max_dna_len=300000, split_hit=20000’. SNAP was trained with the MAKER gff file and a gene prediction with SNAP was carried out inside of MAKER (input: SNAP hmm file, MAKER gff file). SNAP training and the gene prediction were repeated two times.

Gene prediction with BRAKER2 using the primary leaf RNA-Seq mappings, the *Vitis* protein sequences, protein sequences from OrthoDB v10.1 eudicotyledons (Kriventseva et al., 2019) and the UniProt Viridiplantae protein data was performed in mode ‘--etpmode’ that enables prediction with GeneMark-ETP+ v4.61 (Lomsadze et al., 2005; Lomsadze et al., 2014; Bruna et al., 2020) to generate a training set for AUGUSTUS version master_v3.4.0_20201212 (Stanke et al., 2008).

All *ab initio* gene predictions and the data of the initial MAKER gff file were combined with MAKER applying EvidenceModeler v1.1.1 (Haas et al., 2008) with the same parameters as above. tRNAs were predicted with tRNAscan-SE v2.0 (Chan et al., 2021) through MAKER. The gene predictions were quality filtered keeping gene models with AED<0.5 and refined with PASA according to the PASA wiki. Parameters remained the same as used in (Frommer et al., 2022). Transcript variants with identical transcript and CDS sequence of a gene were discarded. The refined gene models were functionally annotated with a BLASTP search against the UniProt/SwissProt DB (release 2020_06) and protein domains identified with InterProScan5 v5.48-83.0 (Jones et al., 2014) and PFAM database v33.1. Genes encoding proteins <50 aa carrying no functional annotation were discarded.

### 4.14 Resistance gene prediction

Genes were classified into RGAs classes based on predicted protein domains with RGAugury v2.1.7 (Li et al., 2016) using the protein sequences as input. As components of the pipeline, the tools ncoil (Lupas et al., 1991), PfamScan v1.6 (Mistry et al., 2007), InterProScan5 v5.45-80.0 with the databases Gene3D-4.2.0, Pfam-33.1, SMART-7.1 and SUPERFAMILY-1.75 and Phobius-1.01 (Kall et al., 2004) were run to identify relevant protein domains. RGAugurys’ BLAST search against its RGAdb to discard protein sequences showing no hit was disabled.

### 4.15 Differential gene expression analysis

The trimmed data were concurrently mapped on the Gf.99-03 haplotype assemblies with HISAT2 v2.2.0 (Kim et al., 2015) disabling softclipping. Mappings were sorted with SAMtools and mapped reads per gene were counted with featureCounts of the package Rsubread v2.4.3 (Liao et al., 2014) utilizing the parameters ‘GTF.featureType=“exon”, GTF.attrType=“gene_id”, allowMultiOverlap=TRUE, largestOverlap=TRUE, countMultiMappingReads=TRUE’. The count table was filtered for genes with a count sum ≤1 (no expression). DEGs were determined with the function DESeq of the package DeSeq2 v1.30.0 (Love et al., 2014) setting ‘ref=”untreated”’ and ‘design= ∼ condition + time + condition:time’. Count data were transformed with the variance stabilizing transformation setting ‘blind=FALSE’. A PCA plot was generated with the function plotPCA. Additionally, the count data were investigated for a significant difference over time using the LRT with the DESeq function (‘test=”LRT”, reduced = ∼ condition + time’). Counts of genes with adjusted p-value<0.05 were plotted with ggplot2 v3.3.5 (Wickham, 2009). Using the R library UpSetR v1.4.0 (Lex et al., 2014; Conway et al., 2017), shared DEGs between the time points were visualized and DEGs were functionally annotated against the KEGG database (Kanehisa et al., 2021) with KAAS v2.1 (Moriya et al., 2007) using the aa sequence, eukaryotes as GENES dataset, BLAST and single-directional best hit.

### 4.16 Design of new markers

To design specific markers within the *Rpv12* locus, the region of the NLR-Cluster of the Gf9918 haplotype was screened for di- tri-, tetra-, penta-, and hexa-repeats using WebSat (Martins et al., 2009). Repeat regions with a maximum of 500 bp flanking sequence were blasted against PN40024 12X.v0, Gf9918 as well as Gf9921. Primer3Plus (Untergasser et al., 2007) was used for testing PCR suitability. Resulting primer pairs were first tested on ‘Tramine’ and ‘Kunbarat’ by PCR and agarose gel electrophoresis and followed by testing for specificity on 18 genotypes (Table S12) using fluorescently labelled forward primers (Table S11, Table S12 & Table S1).

### 4.17 Analysis of *Rpv12* resistance locus

Based on a primer sequence alignment of the *Rpv12* QTL delimiting markers UDV-014 and UDV- 370 and the other *Rpv12* related SSR markers (Venuti et al., 2013), the *Rpv12* locus position was determined on chr14 of Gf9921, Gf9918 and PN40024 12X.v2. As Gf9918 is the genetic source of Gf.99-03s’ *Rpv12* locus, significantly DEGs and RGAs of Gf9918 were considered as potential candidate genes for *Rpv12* mediated *P. viticola* resistance. The *Rpv12* locus was visualized with the R package gggenomes (https://github.com/thackl/gggenomes) allowing blast hits with >85 % query coverage and identity.

Orthologues between the *Rpv12* genes of the haplotypes and PN40024 were determined with OrthoFinder v2.3.11 (Emms and Kelly, 2015; Emms and Kelly, 2019) using the longest protein sequences. For the Gf9918 candidate genes, gene mates positioned in the *Rpv12* locus of the susceptible haplotype Gf9921 were identified based on the orthogroups, their position on the locus and their functional annotation. To further validate the uniqueness of the candidates, the protein sequences were used in a BLASTP search against the Gf.99-03 and PN40024 VCost.v3 proteins. Only protein sequences with a query coverage and identity above or equal to 70 % were further investigated. Protein alignments were built with CLC Genomics Workbench v21.0.1 (gap open cost 5.0, gap extension cost 1.0, free end gap cost).

## 5 Author Contributions

SM did the laboratory work including SSR marker analysis, the *P. viticola* infection experiment and the extraction of RNA at the JKI (Julius Kühn-Institute, Institute for Grapevine Breeding Geilweilerhof, Siebeldingen, Germany). SM performed the pedigree analysis, the analysis of the mycelial growth, designed and tested new SSR markers and created figures and tables for these parts. BF, SM and EZ designed the infection time-course experiment. BF did the bioinformatics and analyses concerning the computation of the genome assembly and gene annotation, the evaluation of haplotype phasing, the DEG analysis and the comparisons of the *Rpv12* locus. Bioinformatic analyses were done at Bielefeld University, Faculty of Biology & Center for Biotechnology (CeBiTec). BF created the figures and tables for the bioinformatic analyses.

DH extracted the DNA for the Illumina sequencing and PV did the library preparation and sequencing of the Illumina data. BH did the DNA extraction, library preparation and sequencing of the PacBio long read data.

DH and BW supervised the work at Bielefeld University and RT and EZ supervised the work at JKI Geilweilerhof.

BF and SM wrote the draft manuscript. EZ and BW edited the manuscript. All authors have read and approved the final manuscript.

## 7 Acknowledgments

We would like to thank the members of the Department of Genetics and Genomics of Plants for their support and active exchange. We thank Abdulkhalek Abdaan for technical assistance, Lucie Gebauer for help with R-Studio and Thomas Proksch for support with Fiji. We acknowledge the financial support of the German Research Foundation (DFG) and the Open Access Publication Fund of Bielefeld University for the article processing charge.

## 9 Supplementary Material

### 9.1 Supplemental methods

**Methods S1. Monitoring necrosis around stomata**

**Methods S2. Phase-separated genome assembly and pseudochromosome construction**

**Methods S3. Correction of mis-assemblies**

**Methods S4. Gene prediction and identification of resistance genes**

### 9.2 Supplemental figures

**Figure S1. Combined staining of inoculated leaf discs of Gf.99-03 with Auramin O and Calcofluor white.**

**Figure S2. All-versus-all dotplots between the pseudochromosomes of the haplotypes and the grapevine reference genotype PN40024.**

**Figure S3. Mis-assembly of the Gf9921 pseudochromosome chr08.**

**Figure S4. Spectra-copy number plot of the Gf.99-03 genome assembly.**

**Figure S5. KEGG pathway ‘plant-pathogen interaction’.**

**Figure S6. Amount of Gf.99-03 genes associated with various RGA classes.**

**Figure S7. Expression over time of the differentially expressed *Rpv12* candidates. Figure S8. Domain comparisons.**

### 9.3 Supplemental tables

**Table S1. SSR marker bioinformatic prediction and analysis.**

**Table S2. SSR marker data to confirm the pedigree of Gf.99-03.**

**Table S3. Read binning of the Gf.99-03 long reads.**

**Table S4. Statistics of the chromosome assignment of the Gf.99-03 haplotypes.**

**Table S5. Pseudochromosome lengths of the Gf.99-03 haplotype assembly.**

**Table S6. k-mer based phasing assessment with Merqury.**

**Table S7. RNA-Seq data of Gf.99-03s’ tissues.**

**Table S8. List of bioinformatically identified potential genes encoding domain structures associated with RGA classes.**

**Table S9. RNA-Seq data of *Plasmopara viticola* infection experiment.**

**Table S10. Plant-pathogen interaction [PATH:ko04626].**

**Table S11. Design of new *Rpv12* correlated SSR markers.**

**Table S12. Testing of new *Rpv12* correlated markers.**

## Data Availability Statement

The SMRT sequence data of Gf.99-03 (ERS11455446), the short sequence data of Gf.99-03 (ERS11455447) and of the parental genotypes Gf.43-21 (ERS11455445) and 65-153-18 (ERS11455444) as well as the haplotype sequences of Gf.99-03 (ERS11872527) have been submitted to the DDBJ/EMBL/GenBank databases under project number PRJEB46083. The haplotype assemblies Gf9918 and Gf9921 at chromosome level with structural and functional annotation are available at DOI 10.4119/unibi/2962800.

The RNA-Seq data of Gf.99-03 tissue leaf (ERS11536190), tendril (ERS11536191), root (ERS11536192) and stem (ERS11536193) have been deposited in DDBJ/EMBL/GenBank under project number PRJEB46084. The RNA-Seq data of the *P. viticola* infection experiment on Gf.99-03 leaves (ERS11536194-ERS11536203) have been submitted to the DDBJ/EMBL/GenBank databases under project number PRJEB46085.

