## Supplemental methods for "Phased grapevine genome sequence of an *Rpv12* carrier for biotechnological exploration of resistance to *Plasmopara viticola*"

### *Supplementary Material*

#### 1 Supplementary Methods

##### 1.1 Methods S1. Monitoring necrosis around stomata

Auramine O (Sigma Aldrich, Merck KGaA, Darmstadt) was used to stain lignin and suberin (Pesquet *et al.*, 2005, Ursache *et al.*, 2018). Following the protocol of (Ursache *et al.*, 2018), samples of `Italia´ and Gf.99-03 were bleached in Ethanol absolute for 15 min in a boiling water bath. Subsequently, the decolourised leaf discs were stained with 0.1 % Auramine O solution in $H_2O$ for 24 hpi at room temperature under 50 rpm shaking. Since *P. viticola* structures at early infection times can only be detected by encysted zoospores, a Calcofluor white staining was carried out concomitantly. For this purpose, 10 µl 1 N KOH and 10 µl Calcofluor white staining solution (CalcofluorWhite M2R 1 g/l, Evans blue 0,5 g/l, Sigma Aldrich, Merck KGaA, Darmstadt) were added to the samples. The samples were evaluated with a Leica microscope equipped with a GFP filter (Leica A, excitation λ=340-380 nm, dichroic mirror at 400 nm, emission of the filter: LP 425).

##### 1.2 Methods S2. Correction of mis-assemblies

A set of 15 potential mis-assemblies or mis-assignments (seven on Gf9921 sequences, eight on Gf9918 sequences) were examined (Figure S2). Two mis-assemblies were confirmed for Gf9921 located on chr08 (Figure S3) and chr18 and three mis-assemblies were validated for Gf9918 located on chr02, chr09 and chr16.

For chr08 of Gf9921, the position of the marker GF08-06 did not comply with its position on PN40024 12X.v2 and the dot plot shows a large translocation. This translocation can also be observed when comparing chr08 of Gf9921 against chr08 of Gf9918 (Figure S3 (C)). As a breakpoint with a clear drop in read coverage to less than four was identified at the start of the translocation in the haplotype-specific read mappings, the contig sequence (4,596,015 bp) was split into two sequences (2,624,111 bp; 1,971,904 bp) and the pseudochromosome sequence was corrected (Figure S3 (B)).

The all-versus-all dot plot between Gf9921 and PN40024 12X.v2 and between Gf9921 and Gf9918 (Figure S2 (A, C)) indicated a misplaced sequence on Gf9921s' chr18 that potentially originated from chr12. The sequence is around 1.5 Mbp large and part of a 8,329,106 bp long contig. Moreover, the marker UDV-120 that was designed for chr12 mapped on the potentially misplaced sequence of chr18. Also, 34 protein RBHs ordered this contig part to chr12, yet the contig had 253 protein RBHs with chr18. The breakpoint with a read coverage drop to three reads was detected on position 6,755,658 bp and the contig split into two sequences (6,755,658 bp and 1,573,448 bp). The larger sequence remained on chr18 and the smaller sequence was assigned to chr12.

The mis-assembly on chr02 of Gf9918 was expressed through a mapping of the three chr06 Markers GF06-12, GF06-13 and GF06-15 on a around 3.5 Mbp sequence region that had no alignment with chr02 of PN4004 12X.v2 or chr02 of Gf9921. The sequence was part of a 10,087,624 bp long contig. The dot plots showed a several Mbp large insertion on chr02 of Gf9918 and the all-versus-all dot plot depicted this sequence to chr06 of PN40024 or chr06 of Gf9921. A coverage drop to five reads was detected and the contig sequence was split into a sequence of 6,625,733 bp and a sequence with 3,461,891 bp. The ~3.5 Mbp large sequence was reassigned to chr06.

Another mis-assembly was found on chr09 of Gf9918 based on the chr04 markers UDV-034, UDV-087, VMC7H3 and GF04-05 and the all-versus-all dot plot with PN40024 and with Gf9921 displayed a large sequence part of chr09 of Gf9918 that matched with chr04 of PN40024 and with chr04 of Gf9921. The ~7 Mbp long sequence part is a subsequence of a 18,073,517 bp large contig and a breakpoint with a coverage drop down to five reads was found at 7,085,569 bp dividing the contig into a 7,085,569 bp and a 10,983,915 bp sequence. The ~7 Mbp large sequence was ordered in reverse orientation to chr04.

The last approved mis-assembly was placed on a 7,971,740 bp long contig of chr16 of Gf9918. The all-versus-all dot plot of Gf9918 against PN40024 and against Gf9921 assigned a part of the contig to chr11 and the chr11 marker GF11-10 and UDV-048 supported the assignment. Furthermore, the contig had a set of 66 protein RBHs with chr16, but the sequence part aligning with chr04 had 23 protein RBHs with chr04. A read coverage drop to three reads was located in the read mappings and the contig split into 3,423,043 bp and 4,548,697 bp. The ~3.4 Mbp sequence was placed on chr11.

#### 2 Supplementary Figures

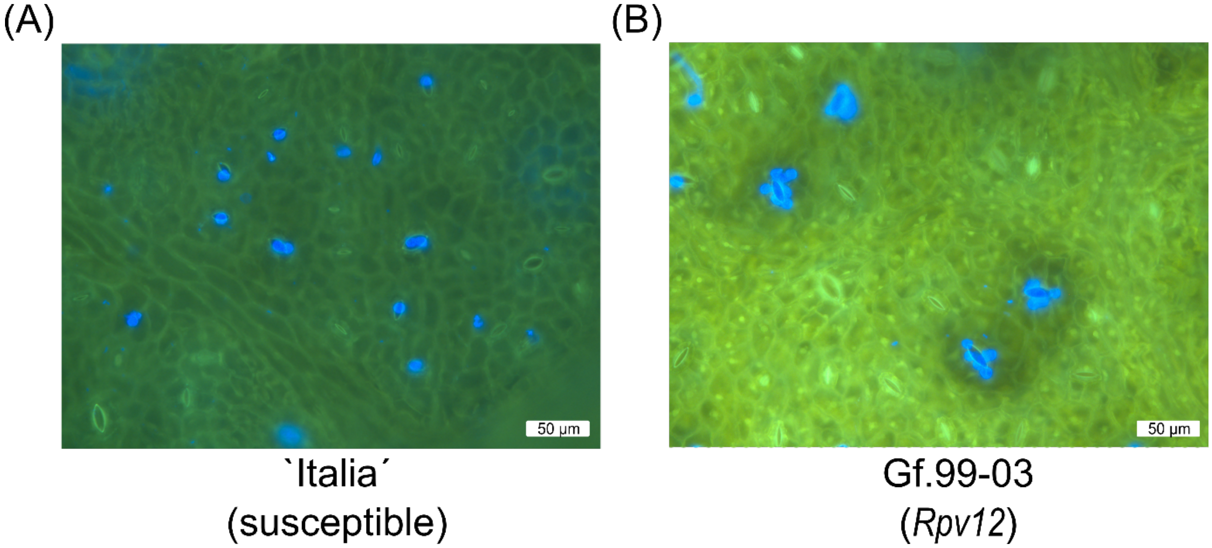


**Figure S1. Combined staining of inoculated leaf discs of Gf.99-03 with Auramin O and Calcofluor white.** The genotypes \`Italia´ and Gf.99-03 were inoculated with zoospores of *P. viticola*. Samples were stained with Auramin O and Calcofluor white at 24 hpi and analyzed using fluorescence microscopy with a GFP-Filter. **(A)** Genotype \`Italia´ shows infected (blue) and uninfected (bright green) stomata. **(B)** Genotype Gf.99-03 shows necrotic regions (brown) around infected stomata (blue).

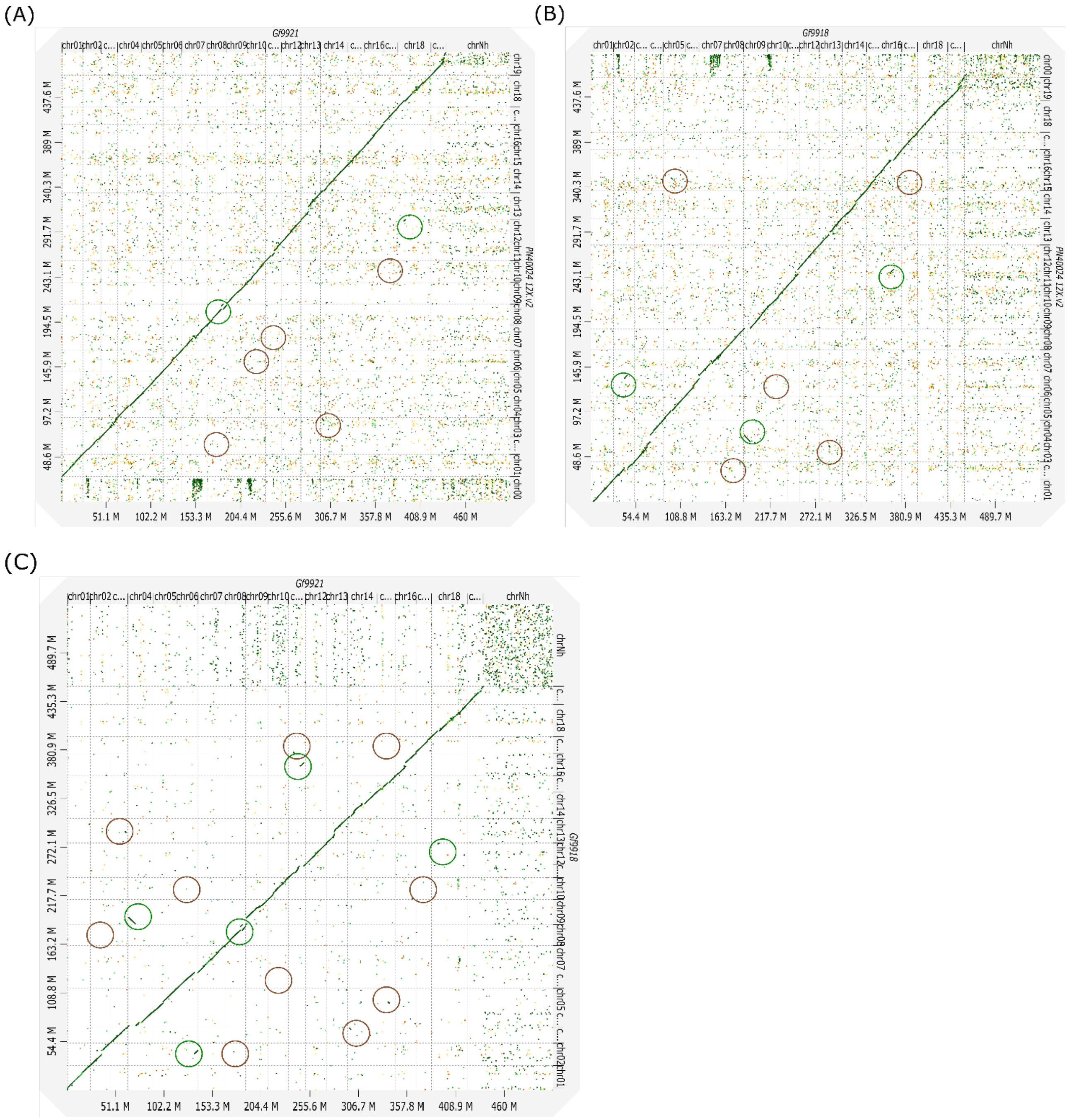


**Figure S2. All-versus-all dotplots between the pseudochromosomes of the haplotypes and the grapevine reference genotype PN40024.** Green circles point on resolved mis-assemblies prior to correction and brown circles point on unresolved potential mis-assemblies. The x- and y-axis are at Megabase pair-scale (M) and comprise the pseudochromosomes. **(A)** Shown is the dotplot of an alignment between Gf9921 and PN40024 12X.v2. **(B)** This dotplot shows the alignment between Gf9918 and PN40024. **(C)** This dotplots shows the alignment between Gf9921 and Gf9918.

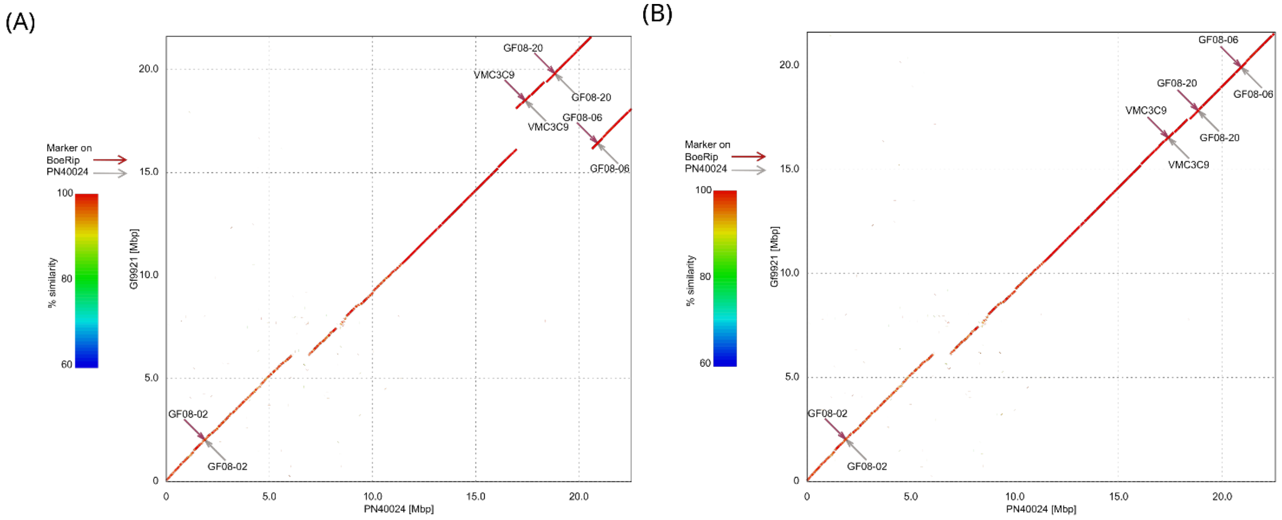


**Figure S3. Mis-assembly of the Gf9921 pseudochromosome chr08.** The dotplots display the alignments between chr08 of the PN20024 12X.v2 assembly (x-axis) and chr08 of the Gf9921 haplotype assembly (y-axis). Magenta arrows denote markers mapping on Gf9921 and grey arrows markers mapping on PN40024. **(A)** This figure shows the alignment prior to mis-assembly correction. **(B)** Figure shows the alignment after the mis-assembly correction.

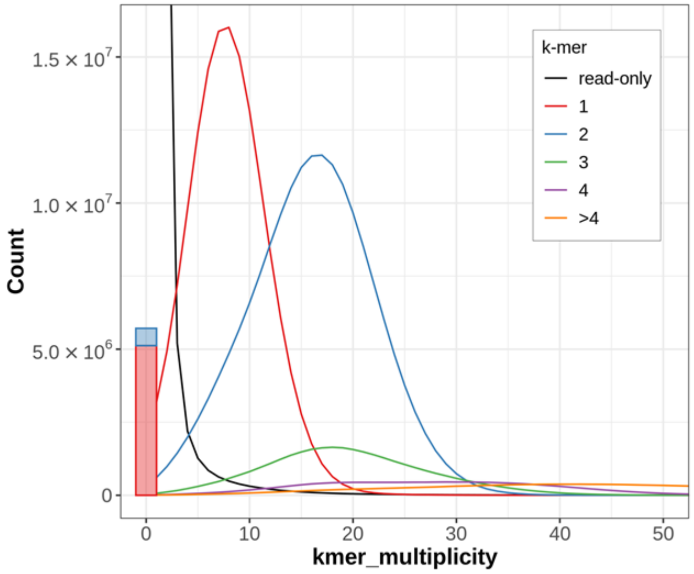


**Figure S4. Spectra-copy number plot of the Gf.99-03 genome assembly.** The figure shows the spectra-copy number plot computed with Merqury. The red peak denotes the 1-copy peak and the blue peak the 2-copy peak. The green peak indicates a small fraction of (artificial) duplications.

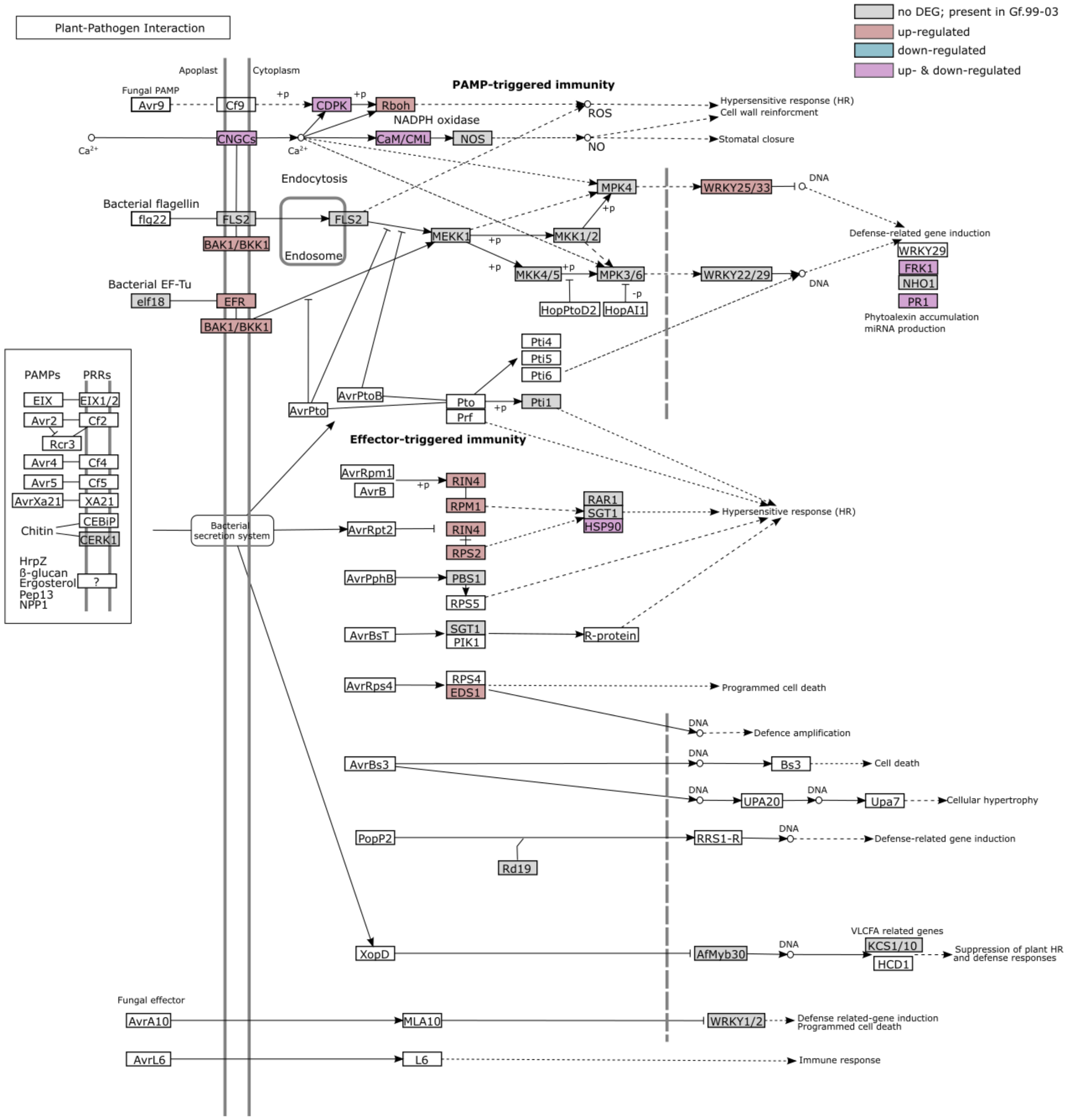


**Figure S5. KEGG pathway 'plant-pathogen interaction'.** The figure shows the KEGG pathway ko04626 'Plant-pathogen interaction'. Proteins highlighted in grey are present in the Gf.99-03 gene annotation. Proteins highlighted in red are in the up-regulated gene set, proteins highlighted in blue are in the down-regulated gene set and highlighted in magenta are both up- and down-regulated. Figure rebuilt from KAAS/KEGG figure 0426 11/18/19 ©Kanehisa Laboratories.

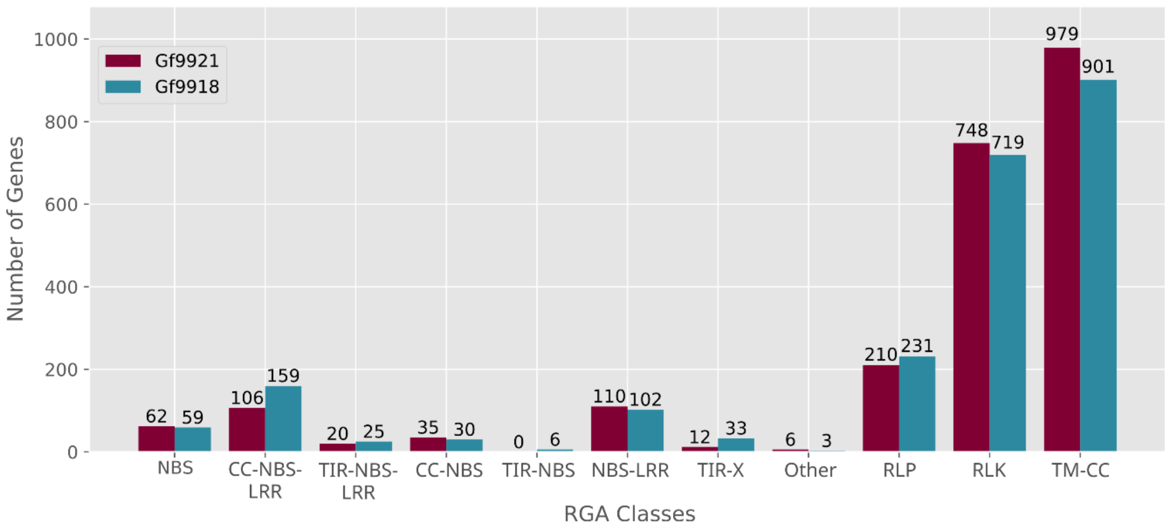


**Figure S6. Amount of Gf.99-03 genes associated with various RGA classes.** Displayed are the number of RGAs for each class and for each haplotype. In total 2,288 RGAs were identified for Gf9921 and 2,268 RGAs were identified for Gf9918. NBS, Nucleotide Binding Site; CC-NBS-LRR with CC for coiled-coil and LRR for leucine-rich repeat; TIR-NBS-LRR with TIR for Toll/Interleukin-1 Receptor like TIR-X with X for unknown site/domain; RLP, Receptor like protein; RLK, Receptor like kinase; TM-CC with TM for Transmembrane.

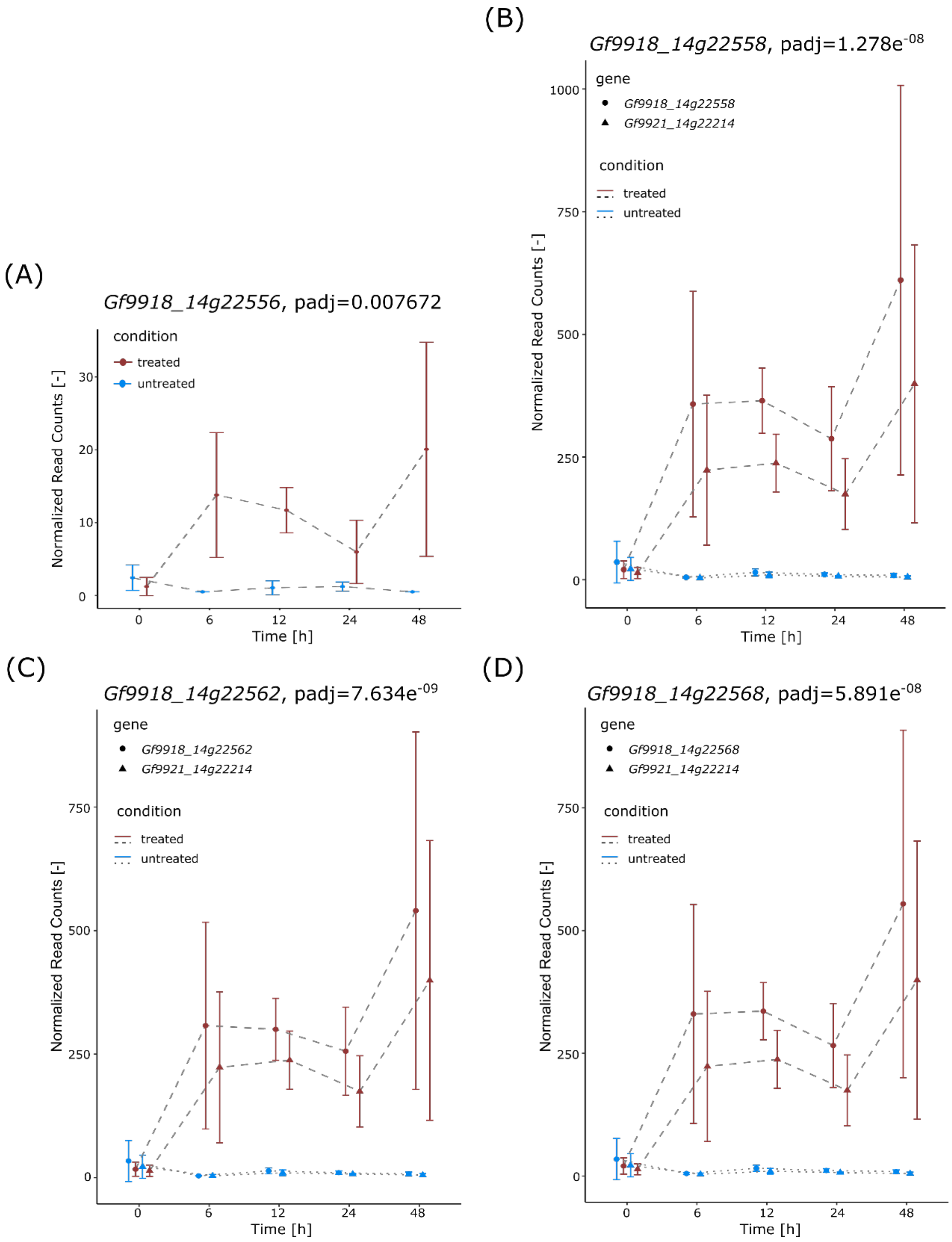


**Figure S7. Expression over time of the differentially expressed *Rpv12* candidates.** The error bars represent the standard deviation. **(A)** Expression course of *Gf9918_14g22556*. **(B)** Expression course of *Gf9918_14g22558* and its gene mate *Gf9921_14g22214*. **(C)** Expression course of *Gf9918_14g22562* and its gene mate *Gf9921_14g22214*. **(D)** Expression course of *Gf9918_14g22568* and its gene mate *Gf9921_14g22214*.

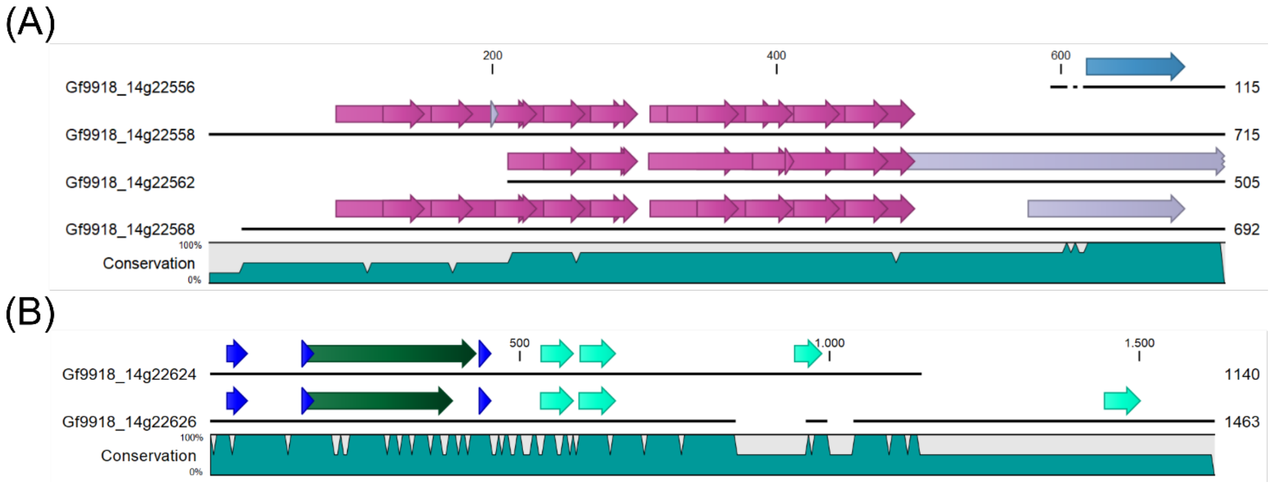


**Figure S8. Domain comparisons. (A)** Comparison of encoded ACD-like Ankyrin-domain carrying proteins. Ankyrin domains are shown in purple, PGG domains are shown in violet and domain of unknown function is shown in blue. Conservation below dyed in petrol shows the matches and deviations between the four proteins. **(B)** Comparison of Gf9918_14g22624 with Gf9918_14g22626. CC-Motifs are shown in blue, NB-ARC domains are shown in green and LRR domains are shown in turquoise. The conservation below dyed in petrol shows the matches and deviations between the two proteins.
